# Senataxin and RNase H2 act redundantly to suppress genome instability during class switch recombination

**DOI:** 10.1101/2021.10.18.464857

**Authors:** Hongchang Zhao, Stella Hartono, Kirtney Mae De Vera, Zheyuan Yu, Krishni Satchi, Tracy Zhao, Lionel Sanz, Frederic Chedin, Jacqueline Barlow

**Author notes:** Corresponding author –.

## Abstract

Class switch recombination generates antibody distinct isotypes critical to a robust adaptive immune system and defects are associated with auto-immune disorders and lymphomagenesis. Transcription is required during class switch to recruit the cytidine deaminase AID—an essential step for the formation of DNA double-strand breaks—and strongly induces the formation of R loops within the immunoglobulin heavy chain locus. However, the impact of R loops on double-strand break formation and repair during class switch recombination remains unclear. Here we report that cells lacking two enzymes involved in R loop removal— Senataxin and RNase H2—exhibit increased R loop formation and genome instability at the immunoglobulin heavy chain locus without impacting class switch recombination efficiency, transcriptional activity, or AID recruitment. Senataxin and RNase H2-deficient cells also exhibit increased insertion mutations at switch junctions, a hallmark of alternative end joining. Importantly, these phenotypes were not observed in cells lacking Senataxin or RNase H2B alone. We propose that Senataxin acts redundantly with RNase H2 to mediate timely R loop removal, promoting efficient repair while suppressing AID-dependent genome instability and insertional mutagenesis.

## Introduction

Class switch recombination (CSR) is a programmed recombination event in mature B cells that generates antibodies of different isotypes, allowing for their interaction with different effector molecules. Successful CSR is a deletional rearrangement catalyzed by the formation of DNA double-strand breaks (DSBs) within the immunoglobulin heavy chain (IgH) switch regions. DSBs are formed by a series of events, initiated by the deamination of cytosine residues in single stranded DNA by activation-induced cytidine deaminase (AID) (Chaudhuri et al., 2003; Muramatsu et al., 2000; Pham, Bransteitter, Petruska, & Goodman, 2003; Ramiro, Stavropoulos, Jankovic, & Nussenzweig, 2003a; Revy et al., 2000). The resulting U:C mismatches are processed by multiple DNA repair pathways producing mutations or single strand DNA (ssDNA) breaks (Stavnezer & Schrader, 2014). In CSR, multiple single-strand breaks on both Watson and Crick DNA strands create double-stranded breaks (DSBs) which initiate recombination between two adjacent but distinct genomic loci. In successful CSR, non-homologous end joining (NHEJ) proteins repair the DSBs by joining the two distal DNA ends, deleting the intervening DNA.

### AID targeting during CSR

AID recruitment to chromatin is a highly regulated act, as off-target AID activity promotes IgH and non-IgH DSBs and translocations associated with carcinogenesis (Ramiro et al., 2004; Robbiani et al., 2008; Robbiani et al., 2009). Transcriptional activity is necessary for the formation of single-strand DNA during CSR, and directly promotes AID recruitment to transcribing switch regions (Chaudhuri et al., 2003; Ramiro et al., 2003a; Zheng et al., 2015). AID interacts with RNA polymerase II cofactors including the transcription factor Spt5 as well as the elongation factor complex PAF1, promoting its recruitment to active switch regions (Pavri et al., 2010; Stanlie, Begum, Akiyama, & Honjo, 2012; Willmann et al., 2012). AID also associates with the ssDNA binding protein RPA (Chaudhuri, Khuong, & Alt, 2004; Yamane et al., 2011). Pre-mRNA splicing also plays a role in AID recruitment, as depletion of the splicing regulator PTBP2 impairs CSR efficiency (Nowak, Matthews, Zheng, & Chaudhuri, 2011). More recently, researchers have shown that AID shows a binding preference for nucleic acid sequences forming G4 quadruplex structures, which are highly enriched in switch sequences (Qiao et al., 2017).

### R loops form at switch regions during CSR

Transcription at the switch region also induces the formation of R loops *in vitro* and *in vivo*—three-stranded nucleotide structures where newly synthesized RNA re-anneals to the DNA template (Aguilera & Garcia-Muse, 2012; Daniels & Lieber, 1995; Reaban & Griffin, 1990; K. Yu, Chedin, Hsieh, Wilson, & Lieber, 2003). While R loops have long been observed at switch regions, their role in CSR remains confounding. The non-template DNA strand of R loops is single-stranded, creating an ideal substrate for AID (Pham et al., 2003; Ramiro, Stavropoulos, Jankovic, & Nussenzweig, 2003b; Sohail, Klapacz, Samaranayake, Ullah, & Bhagwat, 2003). R loop formation in switch regions is sequence-dependent, and positively correlates with AID deamination further suggesting that R loops promote AID recruitment (F. T. Huang et al., 2007). R loops are more stable in GC-rich sequences (Ginno, Lott, Christensen, Korf, & Chedin, 2012; Kuznetsov, Bondarenko, Wongsurawat, Yenamandra, & Jenjaroenpun, 2018; Stolz et al., 2019), and R loop formation in switch region correlates with G density (Zhang, Pannunzio, Hsieh, Yu, & Lieber, 2014). Additionally, G clustering promotes R loop formation in cloned switch regions (Roy, Yu, & Lieber, 2008). Consistent with this, the ATP-dependent RNA helicase DDX1 can promote R loop formation in switch regions, promoting AID recruitment and DSB formation (Ribeiro de Almeida et al., 2018). Yet how R loops are resolved to promote DSB repair is less clear. Studies examining the RNA exosome indicate it promotes AID targeting to both strands of the DNA by helping remove R loops from the template strand, thereby exposing the DNA for deamination (Basu et al., 2011). However multiple studies have used ectopic expression of RNase H1 nuclease to reduce switch region R loops, with contradictory results. One study found that ectopic RNase H1 expression in mouse CH12 cells or primary B cells reduces CSR to IgA or IgG1 respectively, however an earlier study also using CH12 cells observed no effect on CSR to IgA (Parsa et al., 2012; Wiedemann, Peycheva, & Pavri, 2016). Further, transgene-driven expression of murine RNase H1 in primary cells had no effect on CSR to any IgG isotype though somatic hypermutation was increased (Maul et al., 2017). Thus, the enzymes involved in switch region R loop removal and their impact on CSR remain elusive.

Here we use mouse knockout models of two key enzymes implicated in R loop removal, the helicase Senataxin (SETX) and a subunit of the heterotrimeric nuclease RNase H2, to assess the impact of defective R loop removal on DNA double-strand break repair during CSR in primary B lymphocytes (Becherel et al., 2013; Hiller et al., 2012). We found that cells lacking both SETX and RNase H2 activity (*Setx*^*-/-*^*Rnaseh2b*^*f/f*^) exhibit increased R loops specifically at the Sμ switch region, however loss of SETX or RNase H2B alone did not consistently increase R loops in primary B cells. This increase in R loops correlates with enhanced genome instability at the heavy chain locus, as ∼10% of double-deficient B cells contain persistent IgH breaks and translocations. We also observed increased mutations and insertion events in SETX-and RNase H2-deficient B cells by molecular analysis of switch junctions, though there was no defect in CSR efficiency. Taken together, our data suggest that timely R loop removal at switch regions by a SETX/RNase H2 mechanism during class switch recombination suppresses error-prone end-joining and translocation formation at IgH.

## Results

### RNA:DNA hybrids are increased in *Setx*^*-/-*^*Rnaseh2b*^*f/f*^ cells

To determine the effect of R loop metabolism on CSR, we generated mice lacking two enzymes involved in R loop removal, Senataxin and RNase H2. RNase H2 activity is essential for mouse embryogenesis, therefore mice harboring a conditional *Rnaseh2b* allele were crossed to CD19-Cre for B cell-specific gene deletion (*Rnaseh2b*^*f/f*^), then crossed with mice containing a germline deletion of *Setx* (*Setx*^*-/-*^) (**Figure 1A**, (Becherel et al., 2013; Hiller et al., 2012)). Freshly isolated splenic B cells were stimulated with LPS, IL-4 and α-RP105 to induce CSR to IgG1. 72 hours post-stimulation, genomic DNA was isolated and R loop levels were measured by dot blot probed with the RNA:DNA hybrid specific antibody S9.6 (**Figure 1B**). Cells lacking both SETX and RNase H2B showed a 4-fold increase in total R loops, while loss of SETX or RNase H2B alone showed no significant change in R loop levels (**Figure 1B and C**). To determine if R loops were increased within the switch regions, we used DNA:RNA hybrid immunoprecipitation (DRIP) using a monoclonal antibody specific for DNA:RNA hybrids, S9.6. We found a consistent increase in R loop abundance at Sμ specifically in *Setx*^*-/-*^*Rnaseh2b*^*f/f*^ cells, while single mutants exhibited R loop levels similar to WT cells (**Figure 1D**). We also observed DRIP signal at the Sγ1 switch region, however all genotypes exhibited similar levels (**Figure 1D**). We conclude that SETX and RNase H2 act redundantly to remove RNA:DNA hybrids at Sμ.

**Figure 1.**
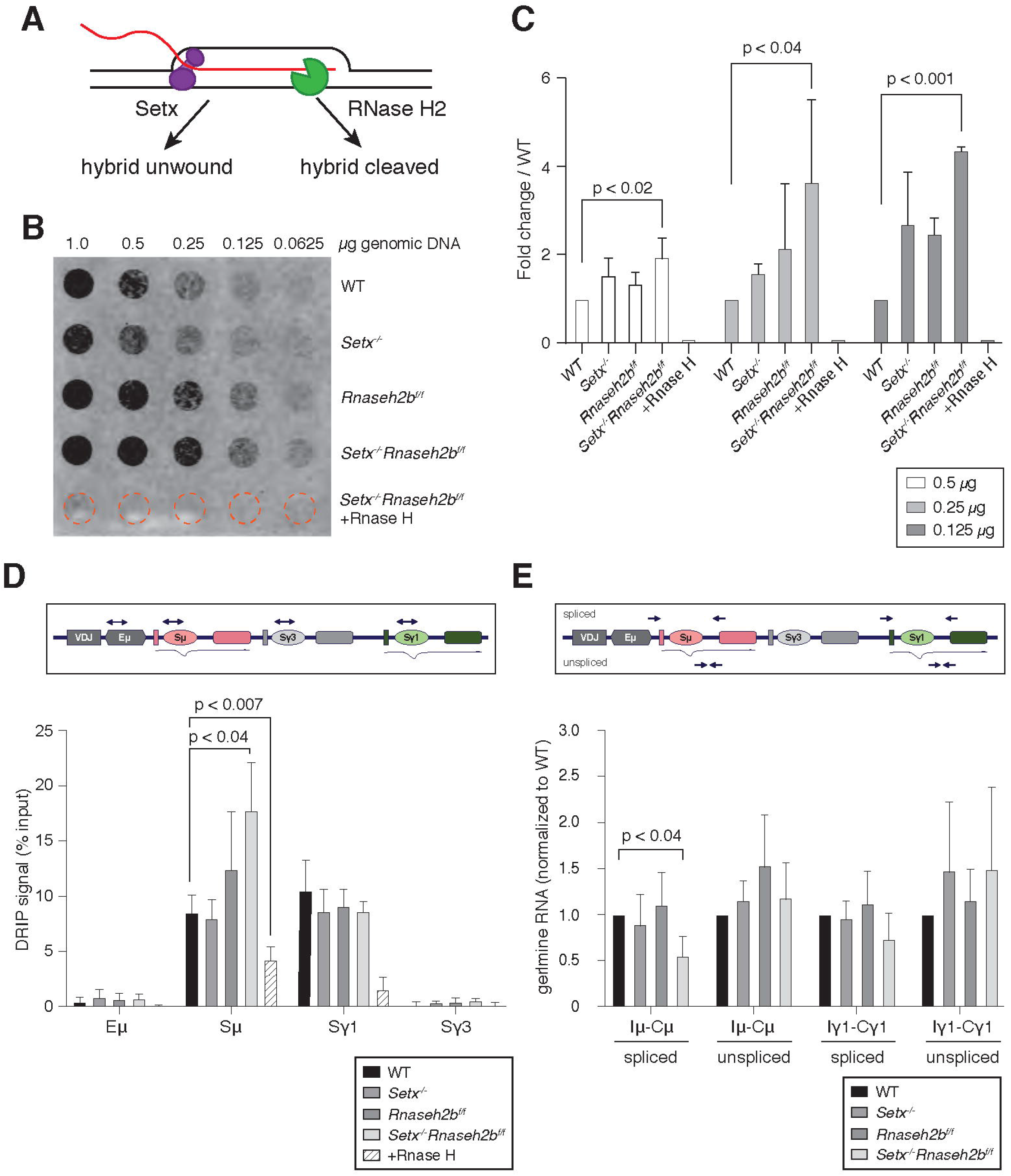
RNA-DNA hybrids are increased in *Setx*^*-/-*^*Rnaseh2b*^*f/f*^ B cells during CSR to IgG1. **A**. Senataxin (SETX) and RNase H2 contribute to RNA:DNA hybrid removal; SETX helicase activity unwinds the nucleotide strands retaining both the RNA and DNA components while RNase H2 cleaves RNA, retaining only the DNA strand. **B**. Dot blot analysis of R loop formation: two-fold serial dilutions of genomic DNA starting at 1 microgram were arrayed on a nitrocellulose membrane and probed using the S9.6 antibody; RNase H treatment of the *Setx*^*-/-*^*Rnaseh2b*^*f/f*^ sample was used as the negative control. **C**. Quantitation of 0.5 micrograms,0.25 micrograms and 0.125 micrograms dot blot from (**B**) using imageJ; values were normalized to WT, set as 1. Figure are expressed as fold-change relative to WT. Error bars are the standard deviation between different experiments; * denotes p < 0.05 comparing different genotypes using multiple T-test (n = 3 mice/genotype). **D**. Diagram of the IgH locus showing the location of PCR products used for ChIP and DRIP assays. DRIP assay was performed with S9.6 antibody using primary B cells after 72 hours of stimulation to IgG1 with LPS/IL-4/α-RP105; RNase H treatment of *Setx*^*-/-*^*Rnaseh2b*^*f/f*^ sample was used as the negative control. Relative enrichment was calculated as ChIP/input and the results were replicated in three independent experiments. Error bars show standard deviation; statistical analysis was performed using one-way ANOVA (n = 3 mice/genotype). **E**. Diagram of the IgH locus showing the location of PCR products for germline transcription under IgG1 stimulation with LPS/IL-4/α-RP105. Real-time RT-PCR analysis for germline transcripts (Ix-Cx) at donor and acceptor switch regions in WT, *Setx*^*-/-*^, *Rnaseh2b*^*f/f*^, and *Setx*^*-/-*^ *Rnaseh2b*^*f/f*^ splenic B lymphocytes cultured for 72 h with LPS/IL-4/α-RP105 stimulation. Expression is normalized to CD79b and is presented as relative to expression in WT cells, set as 1. Error bars show standard deviation; statistical analysis was performed using multiple t-test (n = 4 mice/genotype).

To assess the impact of SETX and RNase H2B loss on an independent locus, we next examined RNA:DNA hybrid signal along the beta-actin gene locus *Actb*. By DRIP-Seq, we found the RNA:DNA hybrid footprint to be largely similar between the four genotypes (**Figure 1-figure supplement 1A**). DRIP-qPCR analysis showed a significant increase in signal at the *Actb* intron 2 locus only in *Setx*^*-/-*^*Rnaseh2b*^*f/f*^ cells, however all other loci examined showed similar RNA:DNA hybrid levels in all genotypes examined (**Figure 1-figure supplement 1B**). Of note, we also observed a trend for reduced DRIP signal in *Setx*^*-/-*^ cells at the promoter and terminator regions. While somewhat surprising, these results are consistent with published reports showing depletion of SETX decreased R loops genome-wide including at the *ACTB* locus (Richard et al., 2020). Together, these results suggest that SETX and RNase H2 impact R loop levels at discrete loci.

### Transcription is not increased in *Setx*^*-/-*^*Rnaseh2b*^*f/f*^ cells

In human cells, the majority of R loop formation positively correlates with gene expression, suggesting that increased transcription can enhance R loop formation (Sanz et al., 2016). To determine if the increased level of R loops at switch regions was due to changes in transcriptional activity, we measured germline transcript levels at switch regions by RT-qPCR 72 hours post-stimulation. In response to LPS/IL-4/α-RP105, we observed similar levels of unspliced germline Sμ and Sγ1 transcripts in all four genotypes (**Figure 1E**). Thus, elevated transcription cannot explain the increased R loop levels at Sμ. Intriguingly, we observed a consistent decrease in spliced Sμ transcripts specifically in *Setx*^*-/-*^*Rnaseh2b*^*f/f*^ cells suggesting that increased or persistent R loop formation may reduce splicing efficiency of germline transcripts specifically at Sμ (**Figure 1E**).

### Cells lacking SETX or RNase H2 are proficient for class switch recombination

R loop formation positively correlates with CSR and is predicted to promote AID recruitment within switch regions, thereby promoting DSB formation (F. T. Huang et al., 2007; K. Yu et al., 2003). Splicing of germline transcripts also correlates with productive CSR, therefore it is possible CSR would be reduced in *Setx*^*-/-*^ *Rnaseh2b*^*f/f*^ cells (Hein et al., 1998; Marchalot et al., 2020). To determine if CSR levels are altered in SETX-and RNaseH2-deficient cells, we measured cell surface expression of IgG1 by flow cytometry. We found that the percent of cells undergoing CSR was similar to WT in *Setx*^*-/-*^, *Rnaseh2b*^*f/f*^ and *Setx*^*-/-*^*Rnaseh2b*^*f/f*^ cells at 72 and 96 hours post-stimulation (**Figure 2A to C**). Thus, neither the observed increase in DNA:RNA hybrid abundance by DRIP nor the reduction in splicing at Sμ significantly impacts CSR to IgG1 in *Setx*^*-/-*^*Rnaseh2b*^*f/f*^ cells. To determine if CSR is proficient to other isotypes, we next measured CSR to IgA and IgG2b by stimulating cells with either LPS/α-RP105/TGF-β/CD40L or LPS/α-RP105/TGF-β, respectively. We found that switching to IgA and IgG2b was also at WT levels in *Setx*^*-/-*^, *Rnaseh2b*^*f/f*^ and *Setx*^*-/-*^*Rnaseh2b*^*f/f*^ cells (**Figure 2D to G**). Together, these results indicate that loss of SETX and RNaseH2 does not impact overall CSR efficiency.

**Figure 2.**
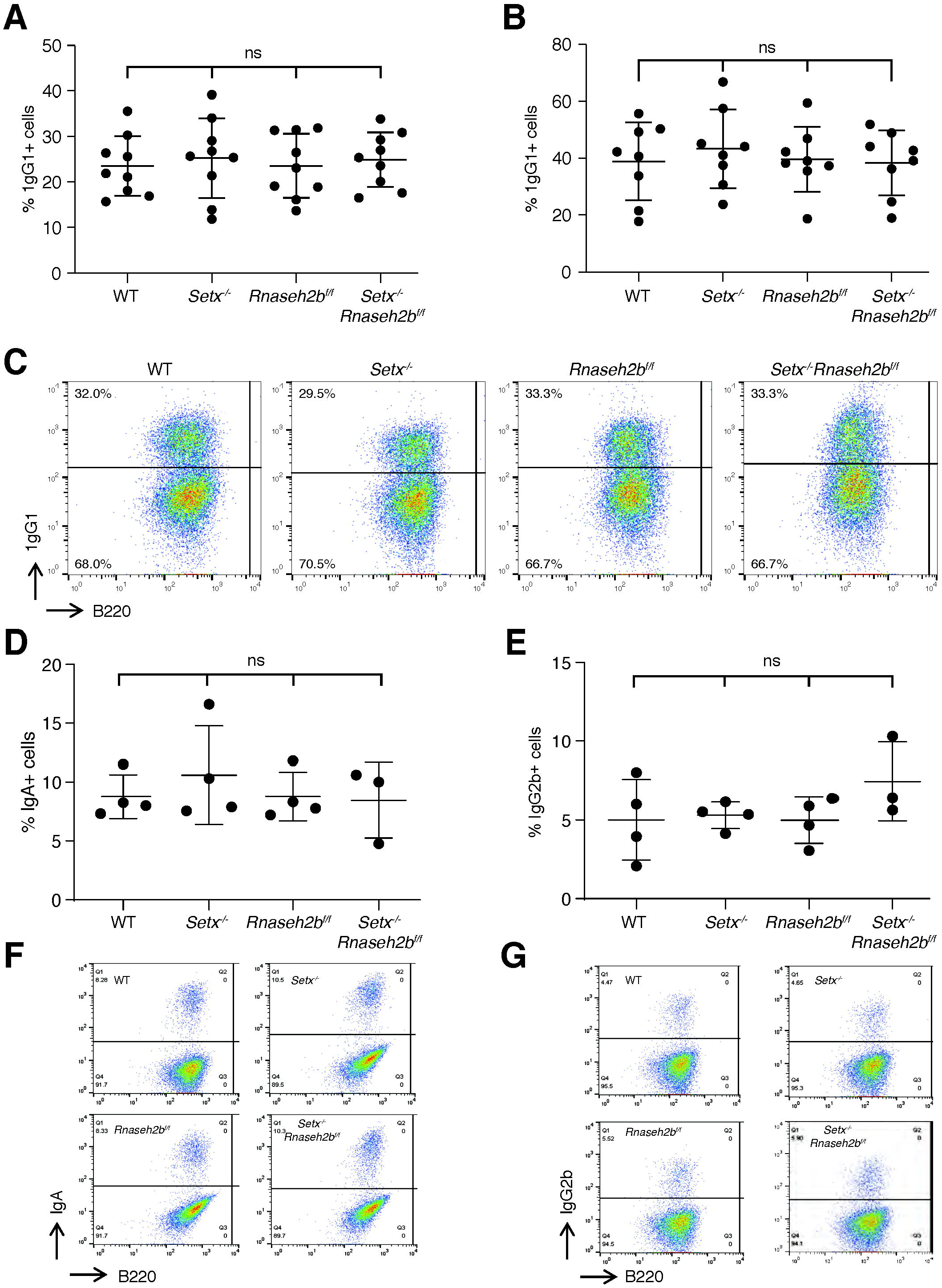
CSR is not reduced in *Setx*^*-/-*^, *Rnaseh2b*^*f/f*^ or *Setx*^*-/-*^*Rnaseh2b*^*f/f*^ B cells. Percentage of cells undergoing CSR to IgG1 72 hours (**A**) and 96 hours (**B**) post-stimulation with LPS/IL-4/α-RP105. **C**. Representative flow cytometry analyses of IgG1^+^ and B220 expression in response to LPS/IL-4/α-RP105. The percentage of IgG1^+^ B cells is indicated at top left. Percentage of cells undergoing CSR to IgA (**D)** or IgG2B (**E**) 72 hours post-stimulation. **F**. Representative flow cytometry analyses of IgA^+^ and B220 expression in response to LPS/α-RP105/TGF-B/CD40L. **G**. Representative flow cytometry analyses of IgG2B^+^ and B220 expression in response to LPS/α-RP105/TGF-B. Horizontal lines in dot plots indicate mean, error bars show standard deviation. Statistical significance versus WT was determined by one-way ANOVA; each dot represents an independent mouse.

### *Setx*^*-/-*^*Rnaseh2b*^*f/f*^ cells have persistent unrepaired DNA damage at IgH

Defects in DNA repair during CSR result in genome instability at IgH visible as persistent DSBs and translocations in miotic chromosome spreads. Defects in R loop removal also correlate with increased DNA damage, therefore we tested if loss of SETX and RNase H2 increased genome instability at IgH. To measure persistent DNA damage and translocations at IgH, we performed fluorescent in situ hybridization (FISH) 72 hours post-stimulation in cells switching to IgG1. Spontaneous damage observed in *Setx*^*-/-*^ and *Rnaseh2b*^*f/f*^ cells was similar to WT levels, however *Setx*^*-/-*^*Rnaseh2b*^*f/f*^ cells harbored significant DNA damage, including chromosome fusions (**Figure 3A**). These results are similar to reports in budding yeast where combined loss of Sen1 and RNase H activity resulted in a synergistic increase in DNA damage (Costantino & Koshland, 2018). No IgH breaks were observed in WT cells and were only found occasionally in *Setx*^*-/-*^ and *Rnaseh2b*^*f/f*^ cells (**Figure 3B**). However, we consistently observed IgH breaks or translocations in ∼10% of *Setx*^*-/-*^*Rnaseh2b*^*f/f*^ cells (**Figure 3B and 3C, n = 4 mice**).

**Figure 3.**
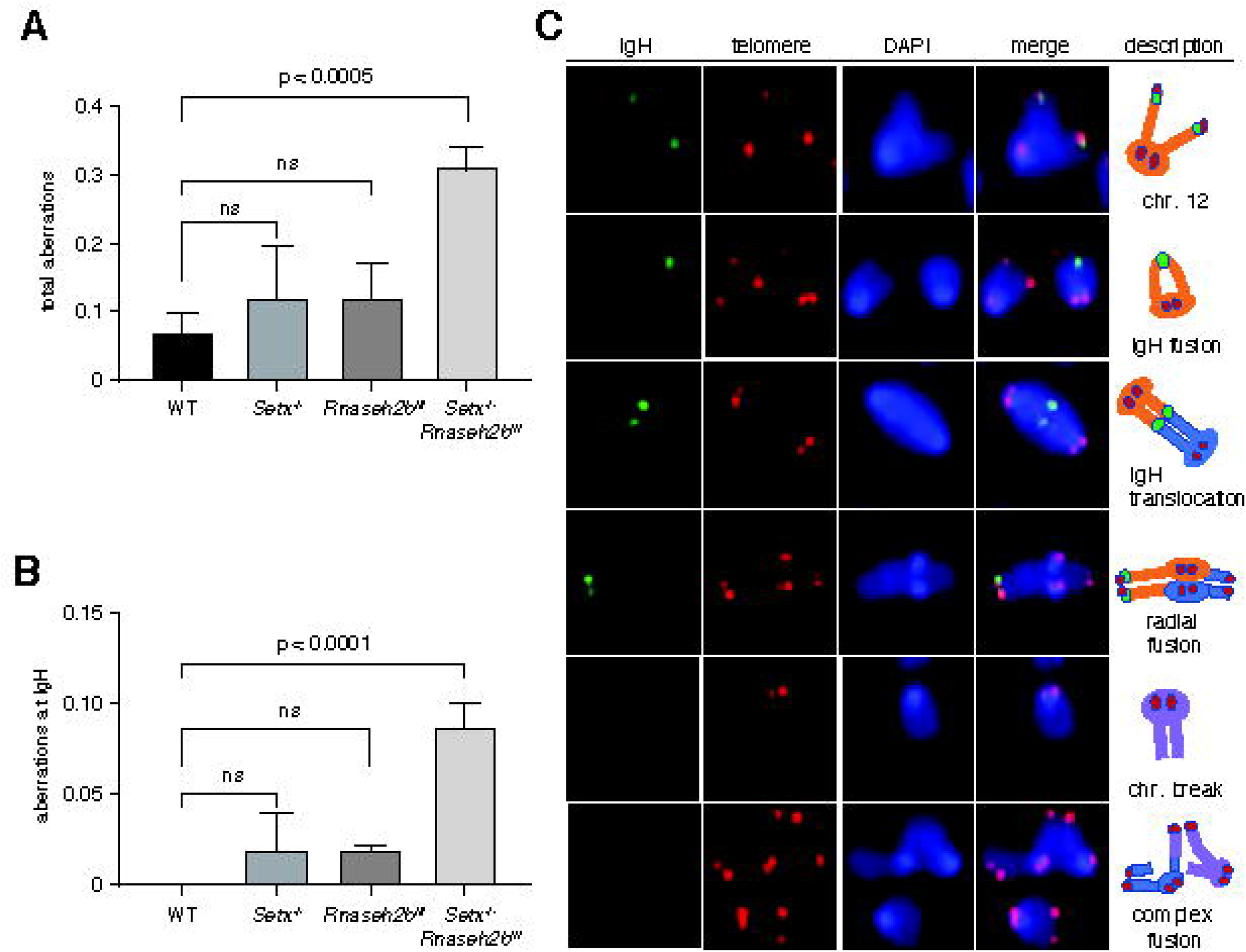
Increased IgH damage is observed in *Setx*^*-/-*^*Rnaseh2b*^*f/f*^ B cells. **A.** Frequency of spontaneous DNA damage in WT, *Setx*^*-/-*^, *Rnaseh2b*^*f/f*^ and *Setx*^*-/-*^*Rnaseh2b*^*f/f*^ cells. **B**. Frequency of spontaneous DNA damage at IgH. **C**. Representative images of the types of rearrangements produced. IgH specific probe visualized in green, Telomere-specific probe visualized in red, DAPI is in blue. All cells were harvested 72 hours post-stimulation to IgG1 with LPS/IL-4/α-RP105. Error bars show standard deviation; statistical significance versus WT was determined by one-way ANOVA (n = 4 independent mice).

### Mis-incorporated ribonucleotides do not contribute to IgH breaks in RNase H2-deficient cells

In addition to cleaving RNA:DNA hybrids, RNase H2 also removes ribonucleotides (RNPs) mis-incorporated into DNA during replication (Hiller et al., 2012; Reijns et al., 2012; Williams & Kunkel, 2014). High levels of genomic RNPs can lead to genome instability, therefore they may also contribute to the persistent DNA breaks observed at IgH. We first measured RNP incorporation by alkaline gel electrophoresis, as DNA enriched for RNPs is sensitive to alkaline hydrolysis leading to single-strand breaks (Nick McElhinny et al., 2010). We found that genomic DNA isolated from *Rnaseh2b*^*f/f*^ and *Setx*^*-/-*^*Rnaseh2b*^*f/f*^ cells is more sensitive to alkaline treatment than DNA from WT or *Setx*^*-/-*^ cells (**Figure 3-figure supplement 1A and 1B**). Importantly, there was no significant increase in RNPs in *Setx*^*-/-*^*Rnaseh2b*^*f/f*^ cells compared to *Rnaseh2b*^*f/f*^ alone. These results indicate that RNPs do not contribute to the total DNA damage observed in *Rnaseh2b*^*f/f*^ and *Setx*^*-/-*^*Rnaseh2b*^*f/f*^ cells, but RNPs do not significantly contribute to the IgH breaks observed specifically in *Setx*^*-/-*^*Rnaseh2b*^*f/f*^ cells. In the absence of RNase H2, the type I topoisomerase Top1 cleaves RNPs from genomic DNA (S. N. Huang, Williams, Arana, Kunkel, & Pommier, 2017; Williams et al., 2013). We hypothesized that excess RNPs would render cells hypersensitive to the Top1 inhibitor camptothecin (CPT), leaving unrepaired breaks in mitosis. Indeed, we found that *Rnaseh2b*^*f/f*^ and *Setx*^*-/-*^*Rnaseh2b*^*f/f*^ cells were sensitive to CPT treatment, showing similar levels of total genome instability (**Figure 3-figure supplement 1C**). Yet CPT treatment did not increase DNA breaks at IgH in any genotype examined (Figure 3B vs. **Figure 3-figure supplement 1D**). We conclude that RNP mis-incorporation does not substantially contribute to IgH breakage in *Setx*^*-/-*^*Rnaseh2b*^*f/f*^ cells even in the presence of exogenous stress.

### Proliferation and cell cycle distribution is normal in cells lacking SETX and RNase H2B

Activated B lymphocytes proliferate extremely rapidly, potentially impacting DNA repair and genome instability (Lyons & Parish, 1994). To determine if cell proliferation is altered, we isolated resting lymphocytes, labeled them with CFSE to track cell division, and stimulated switching to IgG1. After 72 hours, we analyzed CFSE dye dilution and IgG1 expression by flow cytometry. We found that *Rnaseh2b*^*f/f*^ and *Setx*^*-/-*^*Rnaseh2b*^*f/f*^ cells exhibited a modest decrease in cell proliferation (**Figure 3-figure supplement 2A**). CSR frequency correlates with cell division rate (Hodgkin, Lee, & Lyons, 1996); however the CSR frequency was similar in all genotypes examined (**Figure 3-figure supplement 2B**). It is possible that the persistent DSBs observed in *Setx*^*-/-*^ *Rnaseh2b*^*f/f*^ cells trigger DNA damage checkpoint activation, altering cell cycle distribution. Cell cycle phase impacts DSB end resection and repair pathway choice, therefore we analyzed cell cycle profiles by PI staining (Symington, 2016). We found all genotypes had similar fractions of cells in G1 and S phase cells (**Figure 3-figure supplement 2C and 2D**). G2/M cells were modestly increased in *Rnaseh2b*^*f/f*^ and *Setx*^*-/-*^*Rnaseh2b*^*f/f*^ cells, similar to prior reports showing that cells lacking RNase H2 accumulate in G2/M (Hiller et al., 2012) (**Figure 3-figure supplement 2C and 2D**,; p < 0.05, one-way ANOVA). Since *Setx*^*-/-*^ cells did not exhibit an increase in G2/M cells, Thus, we conclude that the increased genome instability at IgH is not due to changes in DNA repair from altered cell cycle distribution.

### RNase H2 activity rescues DNA:RNA hybrid levels in stimulated B cells

It is possible that the increased DNA:RNA hybrids observed in *Setx*^*-/-*^*Rnaseh2b*^*f/f*^ cells arise indirectly due to changes in gene expression or chromatin accessibility earlier in B cell development, as conditional gene deletion using *CD19*^*cre*^ leads to 75-80% deletion in developing pre-B cells in the bone marrow (Rickert, Roes, & Rajewsky, 1997). To determine if RNase H2 activity directly contributes to R loop metabolism at IgH during B cell activation, we expressed FLAG-tagged RNASEH2B in *Setx*^*-/-*^*Rnaseh2b*^*f/f*^ splenic B cells by retroviral infection (**Figure 4A**). Re-expression of FLAG-tagged RNASEH2B suppressed RNP mis-incorporation measured by alkaline gel electrophoresis, indicating the FLAG tag did not substantially interfere with RNase H2 complex formation or its ability to recognize and cleave RNPs covalently attached to DNA (**Figure 4B and 4C**; (Chon et al., 2013)). We also found that RNASEH2B re-expression significantly reduced DNA:RNA hybrid signal at Sµ to levels similar to *Setx*^*-/-*^ cells (**Figure 4D**). Finally, we also observed significant enrichment of RNASEH2B by ChIP at Sγ1 compared to Eµ which does not exhibit high levels of DNA:RNA hybrids (**Figure 4E**; FLAG vs. IgG control). RNASEH2B was also consistently enriched at Sμ relative to IgG, however this result was not significant due to inter-experiment variability (**Figure 4E**). Together, these results indicate that RNase H2 activity contributes to DNA:RNA hybrid removal at IgH during CSR. We were not able to measure SETX binding to IgH as commercially available antibodies did not IP murine SETX (data not shown) and the size of *Setx* precludes expression by retroviral infection, as it encodes for a 2,646 amino acid protein.

**Figure 4.**
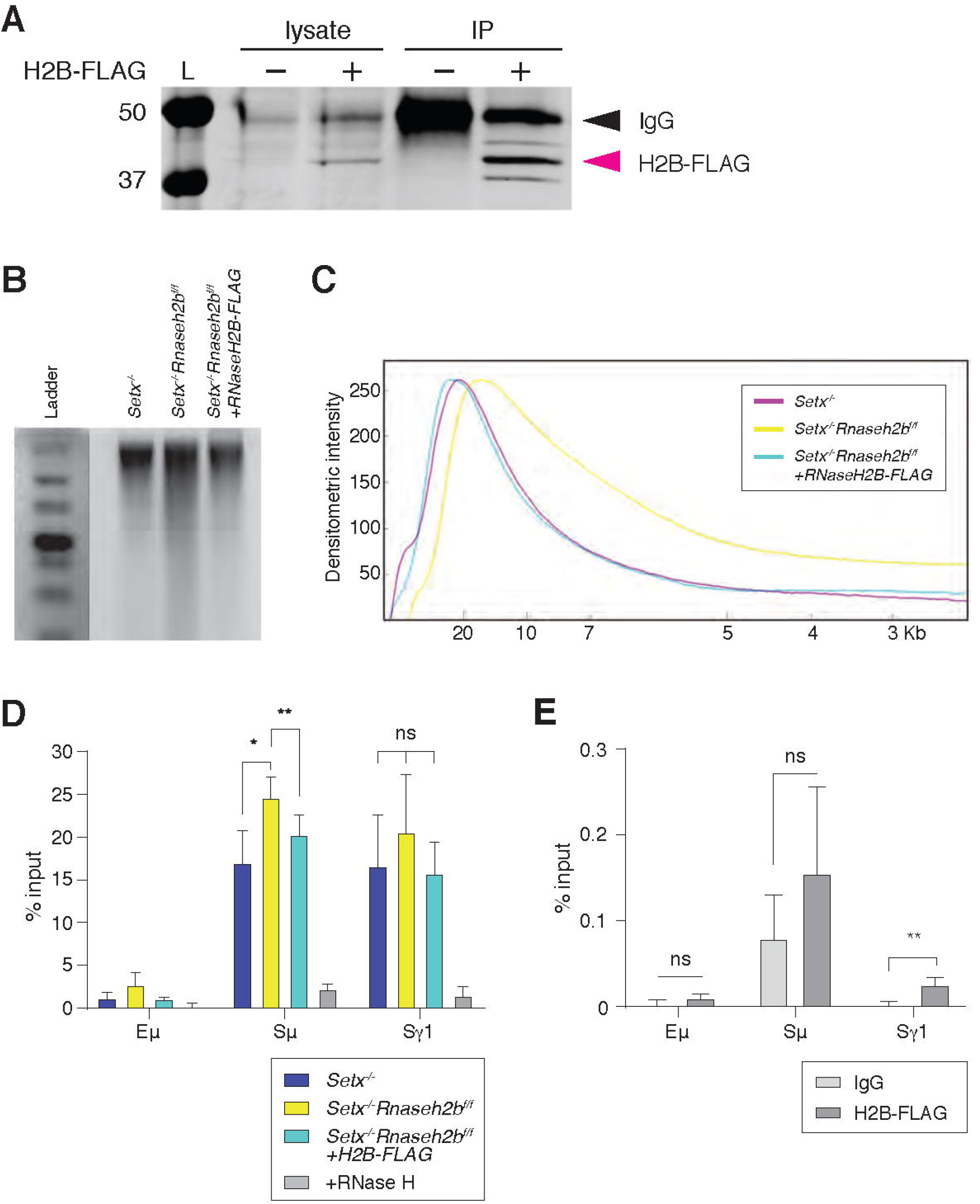
RNase H2 activity rescues DNA:RNA hybrid levels in stimulated B cells. **A**. Total cell lysates were extracted from *Setx*^*-/-*^*Rnaseh2b*^*f/f*^ cells stimulated for 96 hours with LPS/IL-4/α-RP105. Cells were infected with empty vector or retrovirus, expressing FLAG-RNaseH2B, then subjected to immunoprecipitation and immunoblotting with indicated antibodies. **B**. Representative image of alkaline gel from *Setx*^*-/-*^, *Setx*^*-/-*^*Rnaseh2b*^*f/f*^ EV and *Setx*^*-/-*^*Rnaseh2b*^*f/f*^ + FLAG-RNaseH2B cells (n=3 independent mice/genotype). **B**. Densitometry trace of representative alkaline gel in (**A**). **C**. DRIP assay was performed with S9.6 antibody on *Setx*^*-/-*^, *Setx*^*-/-*^*Rnaseh2b*^*f/f*^ and *Setx*^*-/-*^*Rnaseh2b*^*f/f*^ + FLAG-RNaseH2B-expressing cells stimulated with LPS/IL-4/α-RP105 for 96 hours. RNase H treatment of *Setx*^*-/-*^*Rnaseh2b*^*f/f*^ sample was a negative control. Relative enrichment was calculated as ChIP/input and the results were replicated in three independent experiments. Error bars show standard deviation; statistical analysis was performed using one-way ANOVA. **E**. ChIP analysis for FALG-RNaseH2B occupancy in Eμ, Sμ and Sγ regions of primary B cells in response to LPS/IL-4/α-RP105 stimulation. Relative enrichment was calculated as ChIP/input. Error bars show standard deviation. Statistical analysis was performed using student t-test (n = 3 mice/genotype).

### Persistent IgH breaks and translocations are dependent on AID activity

To determine if AID activity is required for the persistent DSBs observed at IgH, we next stimulated cells with α-RP105 alone. Stimulation with α-RP105 induces cell proliferation, however AID expression is minimal compared to stimulation also containing LPS or LPS+IL-4 (**Figure 5A and 5B**; (Callen et al., 2007)). CSR to IgG1 was lower than 1% in all genotypes analyzed, correlating CSR efficiency with AID expression (**Figure 5C**). Total DNA damage levels were similar to LPS/IL-4/α-RP105-stimulated cells, however we did not detect any DSBs or translocations at IgH in α-RP105-stimulated cells for any genotype (**Figure 5D and Supplementary table 1**). These results indicate that the IgH aberrations observed in *Setx*^*-/-*^*Rnaseh2b*^*f/f*^ cells were AID-dependent. Further, only *Setx*^*-/-*^*Rnaseh2b*^*f/f*^ cells consistently accumulate spontaneous unrepaired breaks in the absence of AID expression. It is possible that stimulation with α-RP105 alone alters transcriptional activity and R loop formation within IgH compared to stimulation with LPS/IL-4/α-RP105, altering the potential for R loop-induced IgH breaks. To confirm IgH damage was dependent on AID, we next generated *Aicda*^*-/-*^, *Aicda*^*-/-*^ *Setx*^*-/-*^, *Aicda*^*-/-*^*Rnaseh2b*^*f/f*^ and *Aicda*^*-/-*^*Setx*^*-/-*^*Rnaseh2b*^*f/f*^ mice, stimulated B cells with LPS/IL-4/α-RP105 for 72 hours and performed IgH FISH on metaphase spreads. We observed no IgH breaks in LPS/IL-4/α-RP105-stimulated cells lacking AID, however *Aicda*^*-/-*^*Setx*^*-/-*^*Rnaseh2b*^*f/f*^ B cells had elevated levels of non-IgH damage similar to *Setx*^*-/-*^*Rnaseh2b*^*f/f*^ controls (**Figure 5E**). Further, ∼7% of control *Setx*^*-/-*^*Rnaseh2b*^*f/f*^ cells stimulated side-by-side had IgH breaks, consistent with previous experiments (**Figure 5F and Supplementary table 1**). Together, these results show that the persistent IgH breaks and translocations observed in *Setx*^*-/-*^*Rnaseh2bf/f* cells are AID-dependent.

**Figure 5.**
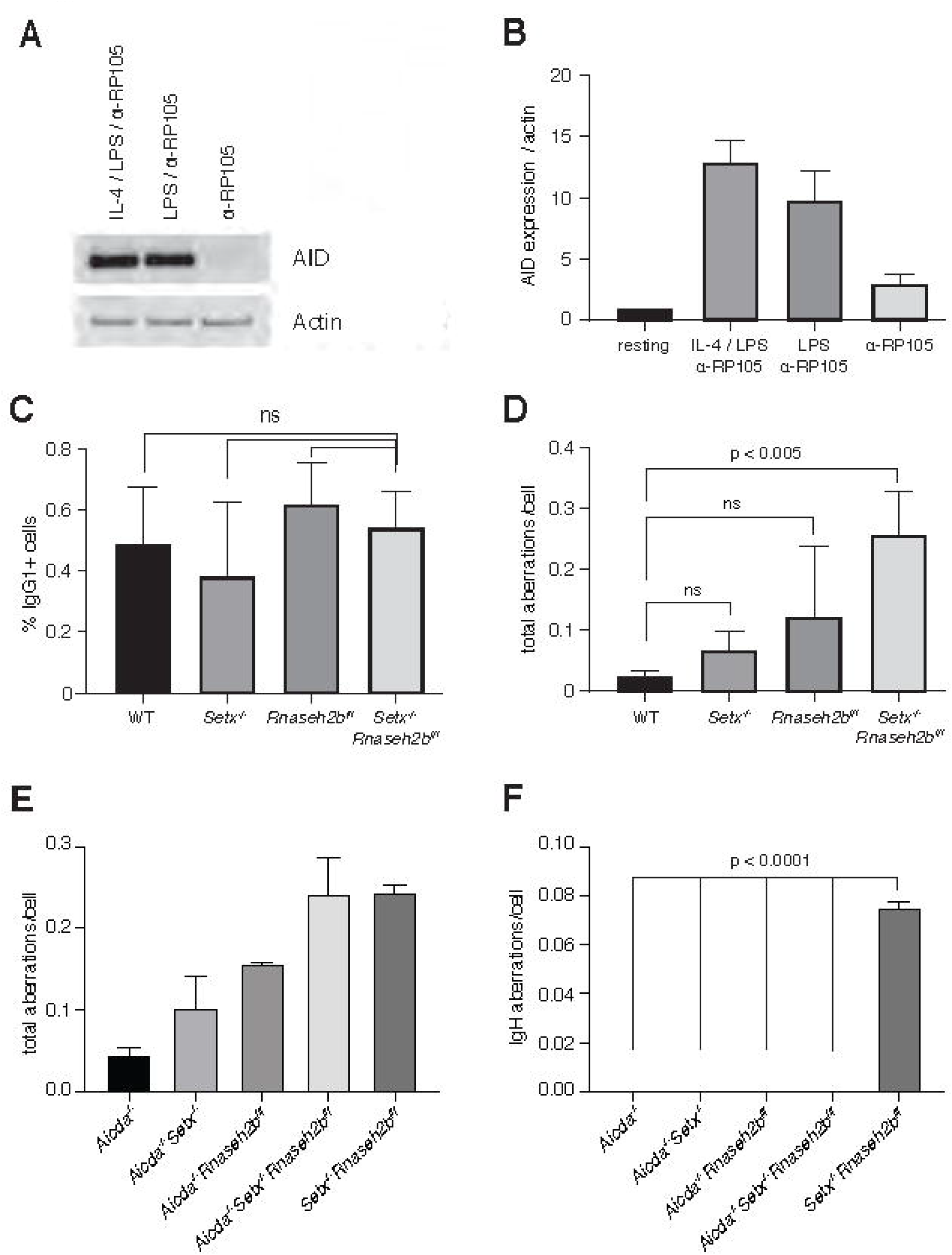
AID activity is required for persistent IgH breaks in *Setx*^*-/-*^*Rnaseh2b*^*f/f*^ B cells. **A**. AID protein levels in WT B cells 72 hours post-stimulation to indicated isotypes (IgG1, LPS/IL-4/α-RP105; IgG3 with LPS/α-RP105; and α-RP105 alone). Actin served as a loading control. **B**. Quantification of AID protein expression relative to Actin for three independent experiments, with AID expression in resting cells set as 1. Error bars show standard deviation. **C**. Percent of cells undergoing CSR to IgG1 in B cells in response to α-RP105 stimulation. Error bars show standard deviation; statistical significance between each genotype was determined by one-way ANOVA(n = 3 mice/genotype). **D**. Frequency of total spontaneous DNA damage under Anti-RP105 treatment in vitro. Error bars show the standard deviation; statistical significance versus WT was determined by one-way ANOVA (n = 3 mice/genotype). **E**. Frequency of total spontaneous DNA damage 72 hours post-stimulation with LPS/IL-4/α-RP105 in *Aicda*^*-/-*^, *Aicda*^*-/-*^*Setx*^*-/-*^, *Aicda*^*-/-*^*Rnaseh2b*^*f/f*^, *Aicda*^*-/-*^*Setx*^*-/-*^ *Rnaseh2b*^*f/f*^ and *Setx*^*-/-*^*Rnaseh2b*^*f/f*^ cells (n = 2 independent mice/genotype). **F**. Frequency of spontaneous IgH damage in *Aicda*^*-/-*^, *Aicda*^*-/-*^*Setx*^*-/-*^, *Aicda*^*-/-*^*Rnaseh2b*^*f/f*^, *Aicda*^*-/-*^*Setx*^*-/-*^*Rnaseh2b*^*f/f*^ and *Setx*^*-/-*^*Rnaseh2b*^*f/f*^ cells 72 hours after stimulation with LPS/IL-4/α-RP105. Error bars show the standard deviation; statistical significance versus WT was determined by one-way ANOVA.

### AID expression and recruitment to switch regions is not altered in SETX or RNase H2-deficient cells

AID overexpression induces high levels of DSBs, increasing class switch recombination efficiency and unrepaired breaks at IgH. To determine if loss of SETX or RNase H2 affected AID expression, we first measured *Aicda* transcript levels. We found that *Aicda* transcripts were similar to WT levels in *Setx*^*-/-*^, *Rnaseh2b*^*f/f*^, and *Setx*^*-/-*^*Rnaseh2b*^*f/f*^ cells indicating that AID gene regulation is not altered (**Figure 5-figure supplement 1A**). AID stability is also regulated during CSR. To determine if protein levels were altered, we next measured protein abundance and found that AID expression was similar in all four genotypes (**Figure 5-figure supplement 1B and 1C**). These results show that the increased damage at IgH is not due to AID overexpression.

Enhanced recruitment of AID to switch regions positively correlates with DSB formation and CSR efficiency. To determine if defective R loop removal alters AID recruitment to switch regions, we performed chromatin immunoprecipitation (ChIP) of AID in WT, *Setx*^*-/-*^, *Rnaseh2b*^*f/f*^, and *Setx*^*-/-*^*Rnaseh2b*^*f/f*^ cells. We found similar levels of AID recruitment at both the Sγ1 and Sµ switch regions in WT, *Setx*^*-/-*^, *Rnaseh2b*^*f/f*^, and *Setx*^*-/-*^ *Rnaseh2b*^*f/f*^ cells by ChIP-qPCR (**Figure 5-figure supplement 1D**). All four genotypes examined showed enrichment at Sγ1 and Sµ compared to *Aicda*^*-/-*^ cells. These results indicate that AID targeting to chromatin is not significantly altered in *Setx*^*-/-*^*Rnaseh2b*^*f/f*^ cells, and enhanced AID targeting to switch regions is unlikely to be the cause of the increased IgH instability.

### RNA polymerase association with switch regions is normal in SETX- and RNase H2-deficient cells

AID physically associates with the transcription factor Spt5, leading to deamination both within and outside switch regions (Pavri et al., 2010; Stanlie et al., 2012). It is possible slower R loop turnover will increase the dwell time of RNA polymerase II (PolII) at R loop-forming genes. To determine if PolII association at switch regions is increased, we performed ChIP of activated PolII (PolII-S5P) in all four genotypes. We found that WT, *Setx*^*-/-*^, *Rnaseh2b*^*f/f*^, and *Setx*^*-/-*^*Rnaseh2b*^*f/f*^ cells all had similar levels of PolII-S5P at both Sμ and Sγ1 switch regions (**Figure 5-figure supplement 1E**). PolII ChIPs showed variability particularly in *Rnaseh2b*^*f/f*^ cells, however IgH breaks were consistently increased *Setx*^*-/-*^*Rnaseh2b*^*f/f*^ cells in every experiment. Thus, we conclude that increased PolII association at switch regions is not the cause of persistent IgH DNA damage observed in *Setx*^*-/-*^*Rnaseh2b*^*f/f*^ cells.

### Switch junctions show elevated insertions in *Setx*^*-/-*^*Rnaseh2b*^*f/f*^ cells

DSBs are generated by creating single strand nicks on the template and non-template DNA strands, potentially creating staggered DSB ends. DSBs with limited overhangs (0-2 nucleotides) are candidates for classical NHEJ (cNHEJ), while breaks containing longer overhangs or mismatches may be repaired by alternative end-joining (alt-EJ). To determine how AID-induced breaks are repaired in SETX and RNase H2-deficient cells, we performed linear amplification-mediated high-throughput genome-wide translocation sequencing (LAM-HTGTS)(Hu et al., 2016). Within Sμ-Sγ1 junctions, we observed a significant decrease in blunt joins specifically in *Setx*^*-/-*^*Rnaseh2b*^*f/f*^ cells (27.3% in WT vs. 17.7% in double-deficient, p = 0.031; **Figure 6A**). In contrast, insertions were significantly increased specifically in *Setx*^*-/-*^*Rnaseh2b*^*f/f*^ cells (21.1% in WT vs 37.7% in double deficient, p = 0.008; **Figure 6A**). Insertion events cells were also longer in *Setx*^*-/-*^*Rnaseh2b*^*f/f*^ than those observed in WT cells, with significant increases in 3 and 4 or more bp insertions (**Figure 6-figure supplement 1C**; p = 0.004 for 3 bp insertions, p = 0.047 for 4 or more bp insertions). *Setx*^*-/-*^ cells also showed increased insertions in Sμ-Sγ1 junctions, however it was not significant (21.1% vs 29.5%, p = 0.059 **Figure 6A**). In addition, junctions with microhomology (MH) were modestly reduced in all mutants, with the largest reduction observed in *Setx*^*-/-*^*Rnaseh2b*^*f/f*^ cells (**Figure 6A**). From these results, we conclude that concomitant loss of SETX and RNase H2 affects repair pathway choice during CSR, increasing error-prone alt-EJ pathways that promote insertions while reducing blunt junctions. Finally, we also observed a 2-fold increase in C>T mismatches in *Setx*^*-/-*^*Rnaseh2b*^*f/f*^ cells (**Figure 6-figure supplement 1D**; p = 0.048). G>A mismatches were also 1.5-fold higher than WT cells, however this was not significantly different (**Figure 6-figure supplement 1D**; p = 0.105).

**Figure 6.**
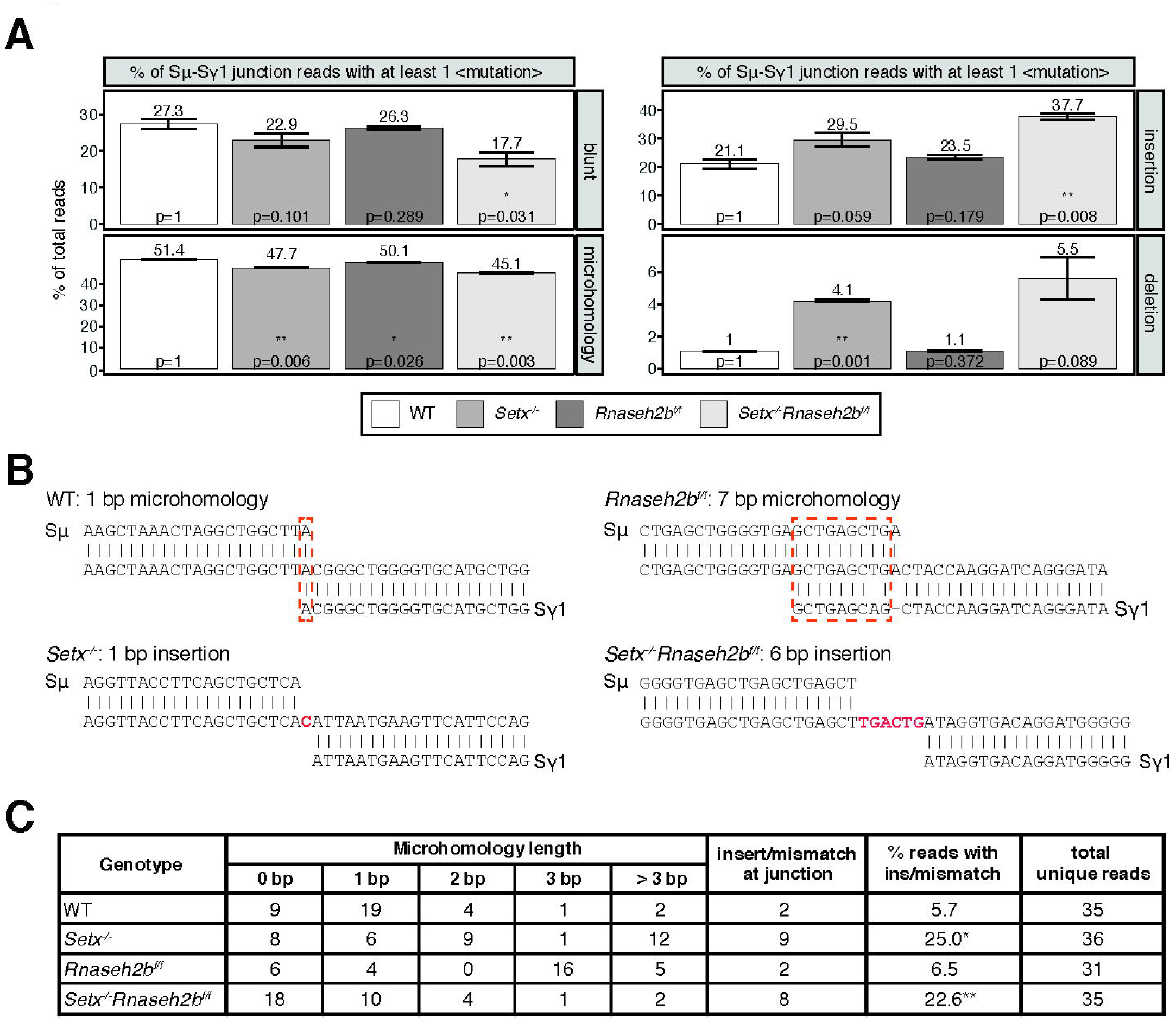
Altered switch junctions in in *Setx*^*-/-*^, *Rnaseh2b*^*f/f*^ and *Setx*^*-/-*^*Rnaseh2b*^*f/f*^ B cells. **A**. LAM-HTGTS analysis showing the percentage of junctions harboring blunt joins, microhomology use, insertions, or deletions at junction sites. LAM-HTGTS uses two technical replicates from genomic DNA isolated from cells 72 hours after LPS/IL-4/α-RP105 stimulation. P values are calculated using student’s t-test. **B**. Representative nucleotide sequences surrounding representative Sμ-Sγ junctions from WT, *Setx*^*-/-*^, *Rnaseh2b*^*f/f*^ and *Setx*^*-/-*^ *Rnaseh2b*^*f/f*^ B cells from Sanger sequencing of cloned junctions. Overlap was determined by identifying the longest region at the switch junction of perfect uninterrupted donor/acceptor identity. Sμ and Sγ1 germline sequences are shown above and below each junction sequence respectively. Regions of microhomology at junctions are boxed with a dashed red line and insertions are in red bold text. Genomic DNA from sequencing experiments was isolated from 2 independent mice for each genotype. **C**. Table with absolute numbers of uniquely mapping cloned switch junctions harboring MH and insertions in WT, *Setx*^*-/-*^, *Rnaseh2b*^*f/f*^ and *Setx*^*-/-*^ *Rnaseh2b*^*f/f*^ B cells 72 hours after stimulation with LPS/IL-4/α-RP105. P-values were calculated using the Chi Square goodness of fit test: *= WT vs *Setx*^*-/-*^ 6.19 × 10^−7^, ** WT vs. *Setx*^*-/-*^*Rnaseh2b*^*f/f*^: 1.25 × 10^−5^).

To further assess junction formation, we cloned and sequenced individual Sµ-Sγ1 switch junctions from all four genotypes. Here *Setx*^*-/-*^*Rnaseh2b*^*f/f*^ cells showed no significant difference in microhomology use from WT cells. This is not surprising as the modest reduction observed in MH use by HTGTS (from 50% to 45%) would be challenging to detect by cloning individual junctions. Similar to HTGTS analyses, *Setx*^*-/-*^*Rnaseh2b*^*f/f*^ and *Setx*^*-/-*^ cells had a significantly increased level of insertions at junction sites compared to WT cells (**Figure 6B and 6C**)—where insertions are defined as junctions containing nucleotides that did not map to either Sµ or Sγ1 (**Figure 6B**)(WT vs *Setx*^*-/-*^ 6.19 × 10^−7^, WT vs. *Setx*^*-/-*^*Rnaseh2b*^*f/f*^: 1.25 × 10^−5^; Chi Square goodness of fit test). Overall, these results support the HTGTS analyses showing an increase in insertion events in *Setx*^*-/-*^*Rnaseh2bf/f* cells indicating an increase in alt-EJ.

## Discussion

In this work, we used combined deletion of *Setx* and *Rnaseh2b* to investigate the role R loops play in CSR. Co-transcriptional R loop formation has emerged as a regulator of CSR involved in targeting AID to the appropriate switch regions for DSB formation. AID recruitment to chromatin is a highly regulated act, as off-target AID activity promotes IgH and non-IgH DSBs and translocations associated with carcinogenesis (Ramiro et al., 2004; Robbiani et al., 2008; Robbiani et al., 2009). However persistent R loops are also sources of replication-associated DNA damage and genome instability (Crossley, Bocek, & Cimprich, 2019; Marnef & Legube, 2021; Prado & Aguilera, 2005; Stork et al., 2016). Here we show that combined loss of SETX and RNase H2 leads to an increase in switch region R loop abundance and IgH damage in the form of unrepaired DSBs and translocations in mitotic cells. While SETX and RNase H2 also have independent functions in DNA repair, loss of either factor alone was not sufficient to significantly increase R loops or IgH instability. The increase in unrepaired breaks was not accompanied by significant alterations in transcriptional activity, splicing or AID recruitment to switch regions indicating that persistent R loops do not impact AID recruitment or deamination. Rather, C>T mutations were increased in *Setx*^*-/-*^*Rnaseh2b*^*f/f*^ cells. Analysis of switch junctions showed an increase in insertion events in *Setx*^*-/-*^*Rnaseh2b*^*f/f*^ cells, indicating that DSB repair by mutagenic alt-EJ was enhanced. From this work, we propose the shared function of SETX and RNase H2 in R loop removal promotes efficient CSR suppresses genome instability during CSR by stimulating efficient NHEJ.

### Separation between CSR efficiency and IgH instability

In contrast to a prior report describing a modest reduction in CSR in SETX-deficient cells, we found that CSR efficiency to IgG1, IgG2B and IgA in *Setx*^*-/-*^ and *Setx*^*-/-*^*Rnaseh2b*^*f/f*^ cells was comparable to WT cells indicating the majority of DSBs created productive junctions leading to cell surface expression (Kazadi et al., 2020). This is consistent with our results which showed no change in germline transcription, RNAP2 association, or AID recruitment. Thus, why do *Setx*^*-/-*^*Rnaseh2b*^*f/f*^ cells have substantially increased IgH breaks without a detectable reduction in CSR? To date, all mouse models exhibiting increased IgH breaks by metaphase spread analysis also exhibit altered CSR efficiency: mice lacking mismatch repair (MMR) factors (MLH1, PMS2, Mbd4), DSB processing factors (CtIP, Exo1), NHEJ proteins (Ku70, Ku80, Xrcc4, Lig4), mediators (53bp1, Rnf8), or kinases (ATM, DNAPKcs) (Alt, Zhang, Meng, Guo, & Schwer, 2013; Bardwell et al., 2004; Boboila et al., 2010; Callen et al., 2007; Casellas et al., 1998; Franco et al., 2008; Grigera, Wuerffel, & Kenter, 2017; Gu et al., 1997; Lee-Theilen, Matthews, Kelly, Zheng, & Chaudhuri, 2011; Li et al., 2010; Lumsden et al., 2004; Manis, Dudley, Kaylor, & Alt, 2002; Reina-San-Martin, Chen, Nussenzweig, & Nussenzweig, 2004; Santos et al., 2010; Schrader, Vardo, & Stavnezer, 2002; Ward et al., 2004; Yan et al., 2007). Alternatively, enhanced AID expression or nuclear localization increases AID-associated damage, however these elevate CSR frequency (Robbiani et al., 2009; Uchimura, Barton, Rada, & Neuberger, 2011). Alt-EJ has been proposed to be a default pathway used when cNHEJ proteins are absent (Bennardo, Cheng, Huang, & Stark, 2008; Boboila et al., 2010; Nussenzweig & Nussenzweig, 2007; Soulas-Sprauel et al., 2007; Stavnezer & Schrader, 2014; Yan et al., 2007), however we report a scenario where unrepaired breaks and mutations at switch junctions are increased without a concomitant reduction in CSR, suggesting that error-prone EJ becomes preferred even when all core cNHEJ factors are present. This is not without precedent, as cells lacking the nuclease Artemis exhibit increased microhomology use at switch junctions without substantially affecting CSR to most isotypes (Du et al., 2008; Rivera-Munoz et al., 2009). Taken together, our results indicate that DSB formation and DNA end joining processes are largely intact in *Setx*^*-/-*^*Rnaseh2b*^*f/f*^ cells, however blunt joins are reduced indicating that DSB repair pathway choice and cNHEJ efficiency are likely impaired.

### DNA structure and AID recruitment

The increase in C>T and G>A point mutations in *Setx*^*-/-*^*Rnaseh2b*^*f/f*^ cells supports a role for R loops in AID recruitment. Crystal structures of AID revealed two nucleotide binding regions—the substrate channel itself and an assistant patch—indicating a preference for branched substrates (Qiao et al., 2017). This raises the possibility that other branched nucleotides could also be AID substrates. Indeed, an RNA-DNA fusion molecule can also bind and be deaminated by AID, raising the notion that R loop “tails” also promote AID recruitment to switch regions (Liu et al., 2021). These structures can work independently or together to enrich AID association along switch regions (**Figure 7**). Of note, R loop tails are flexible, and may cause AID association with template or non-template strands depending on ssDNA availability. R loops in switch regions can also promote the formation of G-quadruplexes in the non-template strand, a structure AID strongly binds (Lim & Hohng, 2020; Qiao et al., 2017). Switch RNAs themselves form G-quadruplexes and AID shows equal binding affinity for RNA and DNA G-quadruplexes (Qiao et al., 2017; Zheng et al., 2015). Thus, R loop stabilization within switch regions may promote the formation of DNA and RNA G-quadruplexes, as well as branched RNA-DNA substrates –all strong substrates for AID binding and activity. However, R loops present a double-edged sword; while their formation may promote AID recruitment and stimulate CSR, their sustained presence potentially perturbs uracil processing, resulting in an increase in C>T and G>A transition mutations during DNA replication.

**Figure 7.**
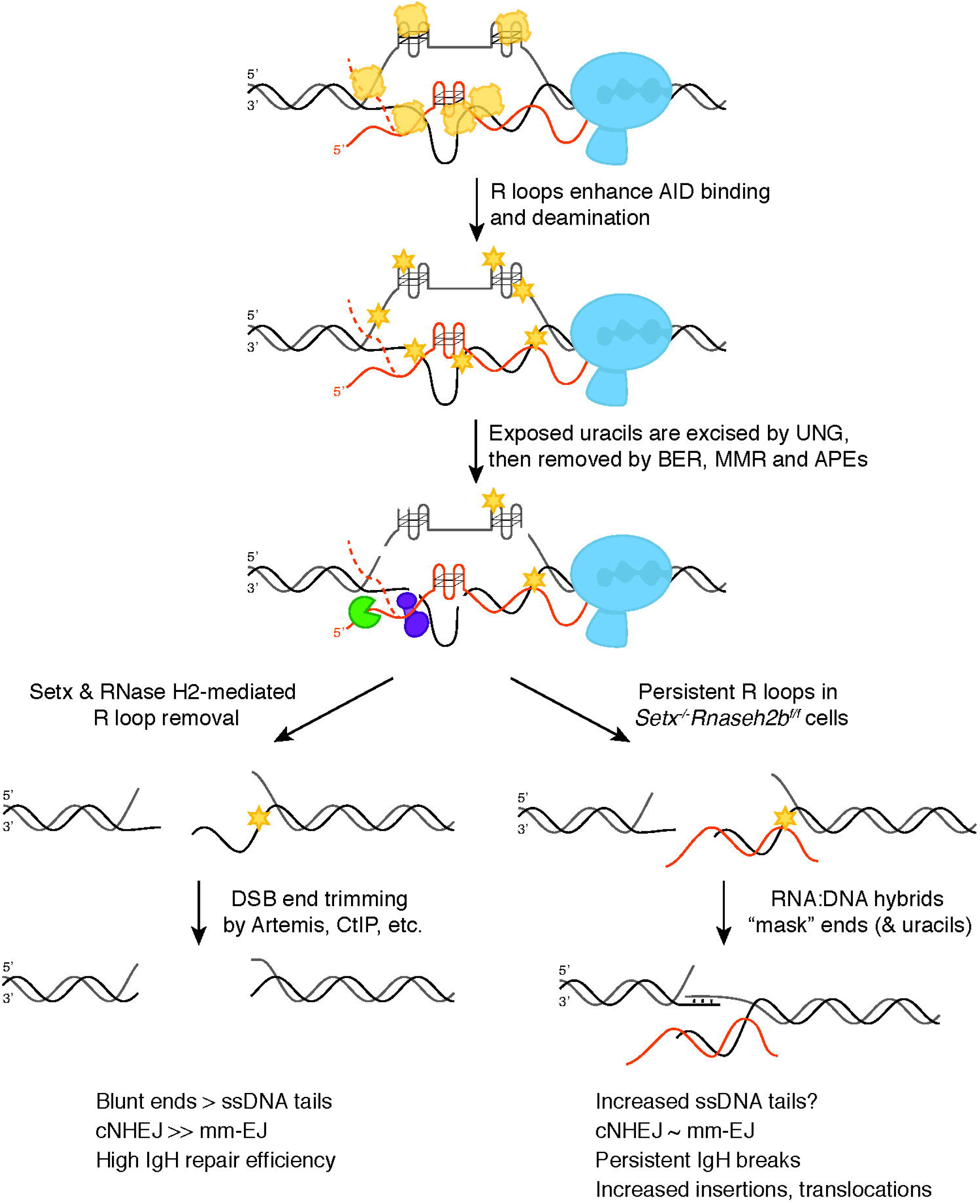
Model for SETX and RNase H2 in promoting efficient CSR. When B cells are stimulated to undergo class switching, PolII-mediated transcription opens duplex DNA at recombining S regions. R loops formed during transcription promote AID binding to ssDNA on the non-template strand first. SETX and RNase H2 then cooperate to remove switch region R loops, exposing ssDNA on the template strand for AID to bind. Extensive AID activity and uracil removal on both strands results in the formation of DSBs with limited ssDNA overhangs at break ends (“blunted” ends) which are predominantly repaired by cNHEJ. When SETX and RNase H2B are absent, R loops forming at S region are not efficiently removed and error-prone EJ is increased. This persistent R loop/RNA:DNA hybrid may affect CSR in two ways. One possible mechanism for increased error-prone EJ is that persistent R loops reduce the extent of ssDNA available for AID binding specifically on the template strand, increasing ssDNA tail length at DSB ends. Alternatively, persistent RNA:DNA hybrids may alter DNA repair protein recruitment to DSB ends, impeding end processing and/or ligation. Both possibilities reduce NHEJ efficiency, but do not affect overall CSR levels as the majority of breaks with long ssDNA tails are repaired by error-prone EJ. However a subset of breaks are not repaired, leading to persistent DSBs that manifest as chromosome breaks and translocations in mitotic spreads.

### Efficient R loop removal promotes cNHEJ and genome integrity during CSR

We propose that R loops at switch regions initially promote AID activity, however their persistence after break formation subsequently interferes with DSB end processing and/or joining, resulting in the IgH instability observed in *Setx*^*-/-*^*Rnaseh2b*^*f/f*^ cells (**Figure 3B and 3C**). This places SETX and RNase H2 downstream of DDX1, an RNA helicase which promotes R loop formation and AID targeting by unwinding G4 quadruplex structures in switch transcripts (Ribeiro de Almeida et al., 2018). Normally, SETX and RNase H2 remove switch R loops along the non-template strand, promoting the formation of DSB ends appropriate for cNHEJ (**Figure 7**). In the absence of SETX and RNase H2, persistent switch region R loops potentially affects CSR in two distinct ways— by changing the type of DSBs created, or by interfering with DSB repair protein recruitment.

In the first possibility, persistent R loops could block AID access to the template strand, potentially increasing the length of 5’ ssDNA tails at DSB ends. This model is supported by a potential role of the RNA exosome in CSR where AID association with RNA exosome components promotes deamination on both template and non-template DNA strands, presumably by removing RNA annealed to the template strand (Basu et al., 2011). However, we found no difference in AID recruitment by ChIP to support this; instead, our observed increase in C>T mutations in switch junctions is an indication of robust AID-mediated deamination. Persistent R loops could also interfere with the recognition and removal of AID-induced uracils, decreasing DSB formation and/or generating DSB ends with 3’ or 5’ ssDNA tails not immediately suitable for cNHEJ. Yet *Setx*^*-/-*^*Rnaseh2b*^*f/f*^ cells are proficient for CSR, demonstrating that processing of deaminated bases successfully produces switch region DSBs visible as persistent IgH breaks in metaphase chromosome spreads. Taken together, we conclude that elevated R loops caused by SETX and RNase H2 loss do not dramatically impede AID recruitment or DSB formation, though DSB end structure may be altered.

Efficient R loop removal may also be necessary after DSB formation. DSBs created during CSR often have staggered ends, requiring further processing for repair by cNHEJ (Stavnezer & Schrader, 2014). Persistent RNA:DNA hybrids at or near DSB ends could slow or block cNHEJ-mediated repair by protecting ssDNA overhangs from processing or by impeding cNHEJ protein binding to DSB ends. Most DSB ends can undergo trimming yielding substrates with short < 3 bp overhangs, suitable for cNHEJ. The Ku70/Ku80 heterodimer has lyase activity, preferentially removing apurinic/apyridimic (AP) nucleotides from the 5’ end of DSBs (5’-dRP) (Roberts et al., 2010). This activity appears restricted to short 5’ overhangs, potentially lessening its interference with HR-mediated repair requiring long 3’ ssDNA tails (Strande, Roberts, Oh, Hendrickson, & Ramsden, 2012; Symington, 2016). Microhomology-mediated end-joining also requires trimming of ssDNA tails prior to ligation (H. H. Chang et al., 2016; So & Martin, 2019). In support of this notion, loss of Artemis increases the frequency of unrepaired IgH breaks and MH use at switch junctions (H. H. Chang et al., 2016; H. H. Y. Chang, Pannunzio, Adachi, & Lieber, 2017; Du et al., 2008; Franco et al., 2008; Rivera-Munoz et al., 2009). Indeed, we found that MH use was reduced 6.3% in *Setx*^*-/-*^*Rnaseh2b*^*f/f*^ cells by LAM-HTGTS (**Figure 6A**). Thus, increased R loops in *Setx*^*-/-*^*Rnaseh2b*^*f/f*^ cells may impede end processing during alt-EJ.

R loop removal also appears important for later steps of HR, thus a role in NHEJ is not unexpected. Reducing R loop removal by SETX depletion does not impair DSB end resection but does slow the recruitment of the strand invasion factor Rad51 (Cohen et al., 2018). Thus, how persistent R loops influence DNA repair likely depends on substrate binding affinity of repair proteins. Early acting sensors as well as helicases and nucleases involved in resection may have evolved to recognize DNA substrates bound by RNA, while proteins catalyzing later steps such as filament formation and D loop formation may not.

Junctions isolated from *Setx*^*-/-*^*Rnaseh2b*^*f/f*^ cells also exhibit a high number of insertion events (**Figure 6A and Figure 6-figure supplement 1C**). Error-prone polymerases such as Pol theta and Pol zeta perform gap-filling around the annealed region in microhomology-mediated EJ, indicating a role for these enzymes in CSR (Mateos-Gomez et al., 2015; Schenten et al., 2009; A. M. Yu & McVey, 2010). Indeed, switch junctions formed in Pol theta-deficient cells notably lack insertions >1 bp, indicating it is required for these events (Yousefzadeh et al., 2014). Thus, R loops may impact the recruitment of specific error-prone polymerases to DSB ends during CSR either directly or indirectly. Further investigation of repair protein recruitment and repair kinetics in *Setx*^*-*^*/-, Rnaseh2b*^*f/f*^, and *Setx*^*-/-*^*Rnaseh2b*^*f/f*^ cells will help delineate how persistent R loops influence DSB end processing and repair pathway choice. Finally, RNA transcripts can also act as templates for recombinational repair mediated Rad52 or microhomology-mediated EJ mediated by Pol theta (Keskin et al., 2014; Mazina, Keskin, Hanamshet, Storici, & Mazin, 2017; McDevitt, Rusanov, Kent, Chandramouly, & Pomerantz, 2018; Storici, Bebenek, Kunkel, Gordenin, & Resnick, 2007). Thus RNAs arising from the switch regions themselves or other transcribed regions may act as templates for the insertions observed at switch junctions.

These two models of how persistent R loops impact CSR are not mutually exclusive, as DSB end structure directly affects repair protein binding affinity and which proteins are necessary for successful repair (H. H. Y. Chang et al., 2017; Serrano-Benitez, Cortes-Ledesma, & Ruiz, 2019; Symington, 2016). We propose that persistent R loops promote the formation of long (> 6 bp) ssDNA tails, increasing the frequency of error-prone EJ at switch joins in *Setx*^*-/-*^, *Rnaseh2b*^*f/f*^, and *Setx*^*-/-*^*Rnaseh2b*^*f/f*^ cells. In the absence of both SETX and RNase H2, RNA:DNA hybrids persist at a subset of DSB ends, interfering with efficient repair by cNHEJ and leading to persistent IgH breaks and translocations observed in mitosis. However, many enzymes—including the RNA exosome, RNase H1, and additional RNA:DNA-specific helicases—can also remove these structures indicating the resolution step of CSR is slowed but not blocked. Indeed, Senataxin is not unique in its ability to unwind RNA:DNA hybrids, therefore additional R loop resolving enzymes may also influence CSR. Multiple helicases have been implicated in R loop removal associated with replication stress including Aquarius, DDX19, and DDX21, among others (Hodroj et al., 2017; Sollier et al., 2014; Song, Hotz-Wagenblatt, Voit, & Grummt, 2017). It will be interesting to determine if these enzymes exhibit similar or distinct effects on CSR.

## Materials and Methods

### Mice

*Setx*^*-/-*^, *Rnaseh2b*^*f/f*^, and *CD19*^*cre*^ mice were previously described and used to generate *Setx*^*-/-*^ *Rnaseh2b*^*f/f*^ *CD19*^*cre*^ mice (Becherel et al., 2013; Hiller et al., 2012; Rickert et al., 1997). For CSR to IgG1, we used a minimum of 10 mice per genotype to reach a power of 0.8 with an expectation to see a 25% difference. A priori power calculations to determine mouse numbers were based on reported CSR data from (Kazadi et al., 2020) and performed using G*power 3.1. Since no difference in CSR was detected for IgG1, subsequent CSR studies were based on similar reports in the literature, using a minimum of n=4 mice per genotype. For molecular analyses, we used n=3 mice to reach a power of 80% for p value calculations estimating differences to be consistently 50% or greater. Primary cells from both male and female mice were used to eliminate sex bias. No difference was observed between sexes. The age of mice was matched in experiments to reduce bias, as this variable is known to alter DNA repair efficiency and CSR. No randomization or blinding was used in mouse experiments. All mouse experiments were performed in accordance with protocols approved by the UC Davis Institutional Animal Care and Use Committee (IACUC protocol #20042).

### B cell stimulation

CD43^-^ resting B cells were isolated using the Dynabeads untouched CD43 mouse B cell isolation kit (Thermo Fisher, 11422D). Isolated B cells were cultured in B cell media (BCM, RPMI-1640 supplemented with 10% fetal calf serum, 1% L-glutamine, 50 IU/ml penicillin/streptomycin, 1% sodium pyruvate, 53 μM 2-mercaptoethanol, 10 mM HEPES). B cells were stimulated with LPS, α-RP105 and interleukin 4 (IL-4) for IgG1, LPS/α-RP105/TGF-B for IgG2b and LPS/α-RP105/TGF-B/CD40L for IgA class switch recombination.

### DRIP analyses (DRIP, DRIP-Seq, dot blot, qPCR)

DRIP and DRIP-Seq were performed as described (Ginno et al., 2012; Sanz et al., 2016). Briefly, after gentle genomic extraction and restriction enzyme fragmentation (Hindlll, Xbal, EcoRI, SspI, BrsGI), 4 micrograms of digested DNA were incubated with 2 micrograms of S9.6 antibody overnight at 4 °C in DRIP buffer (10 mM NaPO4 pH 7.0, 140 mM NaCl, 0.05% Triton X-100). In vitro RNase H digestion was used to generate a negative control. After incubation, antibody-DNA complexes were bound to Protein G Dynabeads and thoroughly washed, DNA was recovered with Chelex-100, the bound fraction was suspended in 0.1 ml 10% Chelex-100 (Bio-Rad), vortexed, boiled for 10 min, and cooled to room temperature. This sample was added to 4 μl of 20 mg/ml Proteinase K followed by incubation at 55°C for 30 min while shaking. Beads were boiled for another 10 min. Sample was centrifuged and supernatant collected. Beads were suspended with 100 μl 2× TE to the beads, vortexed, centrifuged, and supernatants pooled. qPCR was performed with SYBR Select Master Mix (thermo fisher) and analyzed on a Light Cycler 480 (Roche), enrichment calculated by ratio of DRIP/input. For DRIP-Seq analyses, libraries were prepared as described in (Ginno et al., 2012; Sanz et al., 2016).

For dot blot analysis, fragmented genomic DNA was spotted in serial 2-fold dilutions and loaded onto a dot blot device assembled with Whatman paper and nitrocellulose membrane. The membrane was crosslinked in UV-crosslinker 120000uJ/cm2 for 15s, blocked with odyssey TBST blocking buffer, and incubated with the anti-DNA–RNA hybrid S9.6 antibody overnight at 4°C. Goat anti-mouse AlexaFluor 680 secondary antibody (A21057) was used. Quantification on scanned image of blot was performed using ImageJ Lab software.

### Real time quantitative RT-PCR

Total RNA was extracted from stimulated primary B cells with TRIzol (Invitrogen), followed by reverse transcription with ProtoScript II First Strand cDNA Synthesis Kit (NEB) according to the manufacturer’s protocol. qPCR was performed using SYBR Select Master Mix (thermo fisher) and analyzed on a Light Cycler 480 (Roche). Gene of interest/ normalizing gene values ± SD were then normalized to the WT controls; Germline switch region transcription value were normalized to CD79b transcripts before normalized to the WT control. Primers are listed in supplementary table 2.

### Flow cytometry

Primary B cells were washed with PBS and stained with B220-FITC and biotin anti-IgG1 (BD), biotin anti-IgG2b (BioLegend), PE anti-IgA (SouthernBiotech). For biotinylated primary antibodies, cells were then stained with PE-Streptavidin (Beckman Coulter). Data were collected on a BD FACSCanto™ and analyzed using FlowJo software. At least 20,000 events of live lymphoid cells were recorded. For CFSE staining, freshly isolated primary B cells were washed and resuspended in 0.1% BSA/PBS at 1 × 10^7^ cells/ml and labeled with CFSE at a final concentration of 5uM for 10 min at 37°C. CFSE was quenched with ice-cold RPMI 1640 medium containing 10% FCS and washed twice with BCM. Labeled cells were then cultured in BCM and appropriate stimuli for 72 or 96 hours days prior to analysis.

For cell cycle analysis, B cells were harvested 72 hours post-stimulation, washed once with PBS, and resuspended in ice-cold 70% ethanol while slowly vertexing. Cells were fixed overnight, then washed with PBS one time. Cell pellets were resuspended in propidium iodide (PI) staining solution (50 μg/ml PI and 100 units/ml RNase A in PBS), then incubated at room temperature in the dark for 2 hours. Fifty-thousand gated events were collected on a BD FACSCanto™ and analyzed by Flowjo software.

### Metaphase chromosome preparation and fluorescent in situ hybridization (FISH)

The metaphase chromosome preparation and FISH was performed as descripted (Waisertreiger, Popovich, Block, Anderson, & Barlow, 2020). Briefly, Day3 stimulated primary B cells were arrested in metaphase by a 1-h treatment with 0.1 μg/ml demecolcine (Sigma, D1925), treated with 0.075 M KCl, fixed in methanol:acetic acid (3:1), spread onto glass slides and air-dried. FISH were performed on metaphase cells using IgH probe. Prior to hybridization, slides were briefly heated over an open flame, denaturing DNA for IgH detection. Slides were washed in 1 × PBS at room temperature (RT) for 5 min, post-fixed in 1% formaldehyde at RT for 5 min and washed in 1 × PBS at RT for 5 min. Slides were dehydrated in ethanol (75, 85, and 100%) at RT for 2 min each and air-dried. Cells and probes were co-denatured at 75 °C for 3 min and incubated overnight at 37 °C in a humid chamber. Slides were washed post-hybridization in 0.4 × SSC/0.3% NP-40 at 72 °C (2 min), then 2 × SSC/0.1% NP-40 at RT (2 min). Slides were probed with 0.25 μM telomere probe (PNA Bio, F1002) for 1 hour at RT. Slides were then washed in 1 × PBST (1X PBS, 0.5% Triton-X-100) for 5 min at 37 °C. After wash with PBS and dehydrated in ethanol (75, 85, and 100%), Slides were counterstained with Vectashield mounting medium containing DAPI (Vector laboratories Inc., H-1200) before microscopy.

### Microscopy and analysis

B cells were isolated and cultured from a separate mouse for each experiment. A minimum of 50 metaphases were analyzed for each experiment. Metaphases images were acquired using an epifluorescent Nikon microscope with NIS Elements AR4.40.00 software (Nikon). Downstream analysis used ImageJ software (NIH).

### Protein blot and immunoprecipitation

Protein expression was analyzed 72 hours post-stimulation for switching cells unless otherwise indicated. Briefly, 0.5 million cells were suspended with RIPA buffer (50 mM Tris-HCl pH8.0,150 mM NaCl, 2 mM EDTA pH8.0, 1% NP-40, 0.5% Sodium Deoxycholate, 0.1% SDS, Protease Inhibitors) and incubated at 4°C for 30 min, following centrifuge to remove the thick DNA, the whole cell lysis was boiled in protein loading buffer for SDS-PAGE. Immunoblotting was performed with the appropriate primary and secondary antibodies. For FLAG-RNaseH2B immunoprecipitation, total cell lysates were prepared in RIPA buffer and incubated with a desired antibody and appropriate protein A/G-agarose beads at 4°C overnight with gentle agitation. Beads were washed 3 times with lysis buffer, and immunocomplexes were eluted by boiling in SDS sample buffer for 5 min before loading. Anti-AID (Thermo Fisher ZA001) at 1:500 dilution, Anti-β-Actin (Abclonal, AC026) at 1: 100000 dilution, Anti-FLAG (Abclonal, AE005), Goat anti-mouse AlexaFluor 680 secondary antibody (A21057) and Goat anti rabbit AlexaFluor 790 secondary antibodies (A11367) were employed.

### ChIP

CHIP was performed as described (Barlow et al., 2013), Briefly, 1 × 10^7^ stimulated primary B cells were cross-linked with 0.5% formaldehyde for 5 min, then quenched by addition of 125 mM glycine for 5 min at RT. Crosslinked cells were washed with ice-cold PBS three times and then resuspended in ice-cold RIPA buffer (50 mM Tris-HCl pH8.0, 150 mM NaCl,2 mM EDTA pH8.0, 1% NP-40, 0.5% Sodium Deoxycholate, 0.1% SDS, Protease Inhibitors). Chromatin was sheared with a Bioruptor (Diagenode) ultrasonicator to the size range between 200-1000 bp. Samples were centrifuged and supernatant collected, 1% percent of lysate as whole cell DNA input. Antibody-coupled Dynabeads Protein G (Thermo Fisher) were used for immunoprecipitations performed overnight at 4°C. Anti-AID (Thermo Fisher ZA001), Anti-Pol ll Serine5 Phospho (4H8, AB5408) and IgG (AB37355) was used for immunoprecipitation. Beads were washed once in each of the following buffers for 10 min at 4°C: Low salt buffer (0.1% SDS 1% Triton X-100 2 mM EDTA 20 mM Tris-HCl pH 8.0 150 mM NaCl), high salt buffer (0.1% SDS 1% Triton X-100 2 mM EDTA 20 mM Tris-HCl pH 8.0 500 mM NaCl), LiCL buffer (0.25 M LiCl 1% NP-40 1% Sodium Deoxycholate 1 mM EDTA 10 mM Tris-HCl pH 8.0), and TE buffer (50 mM Tris, pH 8.0, 10 mM EDTA). DNA was recovered with Chelex-100 and analysis by qPCR. Data were analyzed using the comparative CT method. Fold enrichment was calculated as ChIP/Input. Primers used for qPCR are listed in supplementary table 2.

### Alkaline gel electrophoresis

For alkaline gel electrophoresis, genomic DNA was extracted from day 3 LPS/IL-4/α-RP105-stimulated B cells. 2ug of genomic DNA was incubated in 0.3 M NaOH for 2 hours at 55°C and separated on an 0.9% agarose gel (50 mM NaOH, 1 mM EDTA) as previously described (Mcdonell, Simon, & Studier, 1977). Gels were neutralized with neutralizing solution (1M Tris-HCL, 1.5M NaCL) and stained with ethidium bromide prior to imaging. Densitometry was analyzed with ImageJ software (NIH).

### Retroviral preparation and B cell infection

Viral infection was performed as described (Waisertreiger et al., 2020). After verification of infection efficiency by flow cytometry, cells were harvested for genomic DNA extraction, DRIP, immunoprecipitation-WB, and FISH.

### Junction analysis

Sμ-Sγ1 switch junctions were amplified using published primers (Zan et al., 2017). Briefly, genomic DNA was prepared from 72 h LPS/IL-4/α-RP105-stimulated B cells. PCR products were cloned using pGEM®-TA cloning kit (Promega) and sequenced with T7/SP6 universal primers. Sequence analysis was performed using the Snap gene software.

### LAM-HTGTS library preparation and analysis

Genomic DNA was isolated from WT, *Setx*^*-/-*^, *Rnaseh2b*^*f/f*^, and *Setx*^*-/-*^*Rnaseh2b*^*f/f*^ cells 72 hours after stimulation to switch to IgG1 with 72 h LPS/IL-4/α-RP105. Linear amplification-mediated high-throughput genome-wide translocation sequencing (LAM-HTGTS) libraries were prepared from genomic DNA as previously described (Hu et al., 2016; Yin, Liu, Liu, & Hu, 2019; Yin, Liu, Liu, Wu, et al., 2019). Sequences were aligned to custom genomes, substituting the mm9 sequence from (114, 494, 415–114, 666, 816) with sequence from the NG_005838.1 (Genbank accession no. NG_005838.1) C57BL/6 IgH sequence on chr 12 (to 11,172 to 183,818), or sequence from the AJ851868.3 (GenBank accession no AJ851868.3) 129 IgH sequence (1,415,966– 1,592,715). Sequences were analyzed as detailed in (Crowe et al., 2018; Hu et al., 2016) with the following adjustments. Junctions were analyzed by comparing sequences to both C57BL/6 and S129 backgrounds. A consensus genome was generated between C57BL/6 and S129, and if reads fell into either category they were deemed not mutated—this eliminated over-estimation of insertions, deletions, and mismatch mutations. Of note, no junctions aligned to S129 on one side and C57BL/6. Deletions are defined as regions missing nucleotides adjacent to prey-break site but having 100% homology in flanking regions. Insertions are defined as regions containing nucleotides that map to neither the bait nor the prey-break site. Microhomologies (MHs) are defined as regions of 100% homology between the bait and the prey-break site. Blunt junctions are considered to have no MHs or insertions. Additional scripts used to analysis can be found in github (https://github.com/srhartono/TCseqplus).

## Acknowledgements

This work was supported by research funding from the National Cancer Institute K22CA188106 and National Institute for General Medical Studies grants 1R01 GM134537 (JHB) and R35 GM139549 (FC). The sequencing was performed by the DNA Technologies and Expression Analysis Cores at the University of California Davis Genome Center, supported by National Institutes of Health Shared Instrumentation Grant **(S10 OD010786-01)**. This study utilized the University of California Davis Cancer Center Flow Cytometry core partially supported by National Institute of Health grant S100D018223. Thanks to Drs. Klaus Rajewsky, Martin Lavin, and Axel Roers for mouse models. We would like to thank Dr. Commodore St Germain and all members of the Barlow and Chedin labs for helpful discussions and suggestions, and Jack McTiernan for assistance with figure design.

## Author contributions

JHB and HZ conceived the study and wrote the manuscript with input from all authors. HZ performed all the experiments. KMDV, KS, and TZ assisted with FISH experiments. ZY performed biostatistics analyses on sanger sequencing data. LS performed DRIP-Seq experiments and SH analyzed LAM-HTGTS data. JB designed the study, supervised the research, and wrote the manuscript.

## Figure legends

**Figure 1-figure supplement 1.**
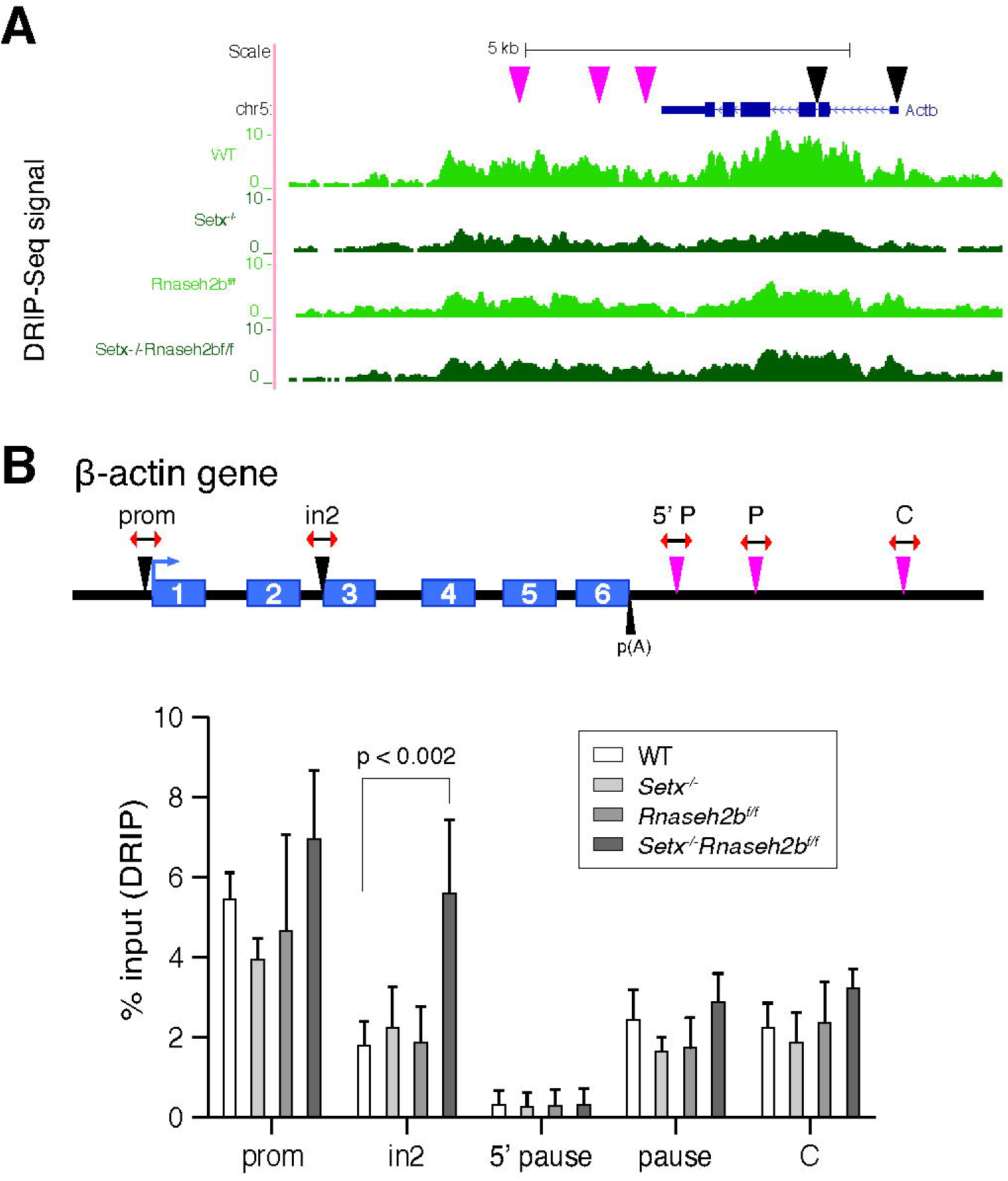
DNA:RNA hybrid formation at the *Actb* locus. **A**. Representative UCSC genome browser screenshot of DRIP-Seq signal in WT, *Setx*^*-/-*^, *Rnaseh2b*^*f/f*^ and *Setx*^*-/-*^ *Rnaseh2b*^*f/f*^ cells spanning 10 kb of the *Actb* locus (n=2 mice/genotype). Locations of primers used in DRIP-qPCR shown in (**B**) are marked with black (gene body) and pink (3’UTR) triangles. **B**. DRIP-qPCR along *Actb* locus in mouse primary B cells. Error bars show the standard deviation from three independent experiments. Statistical analyses were performed using one-way ANOVA, n = 3 mice/genotype.

**Figure 3-figure supplement 1.**
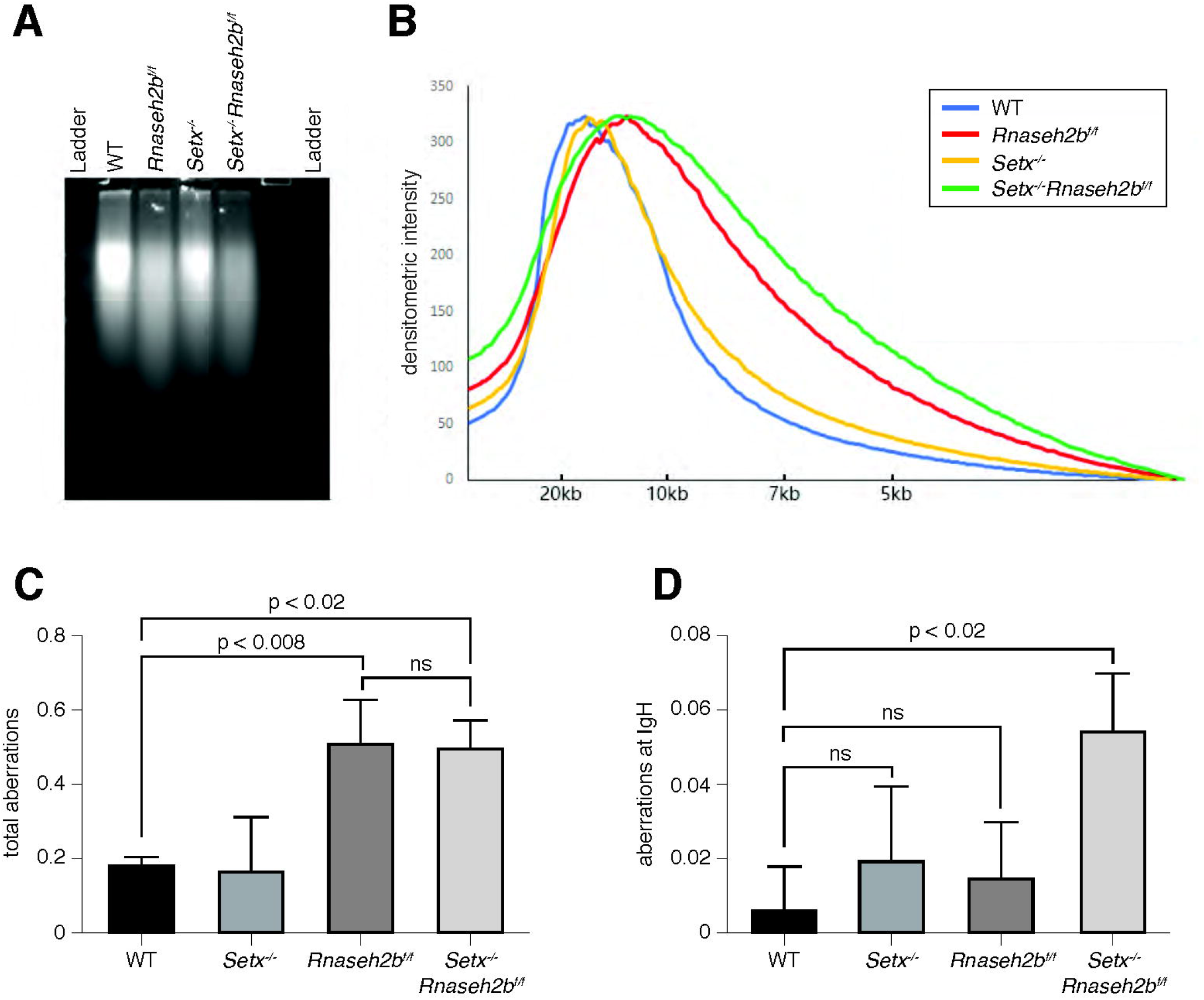
RNP incorporation and camptothecin sensitivity in cells lacking SETX and RNase H2. **A**. Representative image of alkaline gel of genomic DNA (n=3 independent mice/genotype). **B**. Densitometry trace of representative alkaline gel shown in (**A**) with ImageJ software. **C**. Frequency of DNA damage in WT, *Setx*^*-/-*^, *Rnaseh2b*^*f/f*^ and *Setx*^*-/-*^*Rnaseh2b*^*f/f*^ cells after exposure to 5 nM camptothecin (CPT) for 20 hours. **D**. Frequency of DNA damage at IgH in CPT-treated cells. All cells were harvested 72 hours post-stimulation to IgG1 with LPS/IL-4/α-RP105. Statistical significance versus WT was determined by one-way ANOVA (n = 3 independent mice/genotype).

**Figure 3-figure supplement 2.**
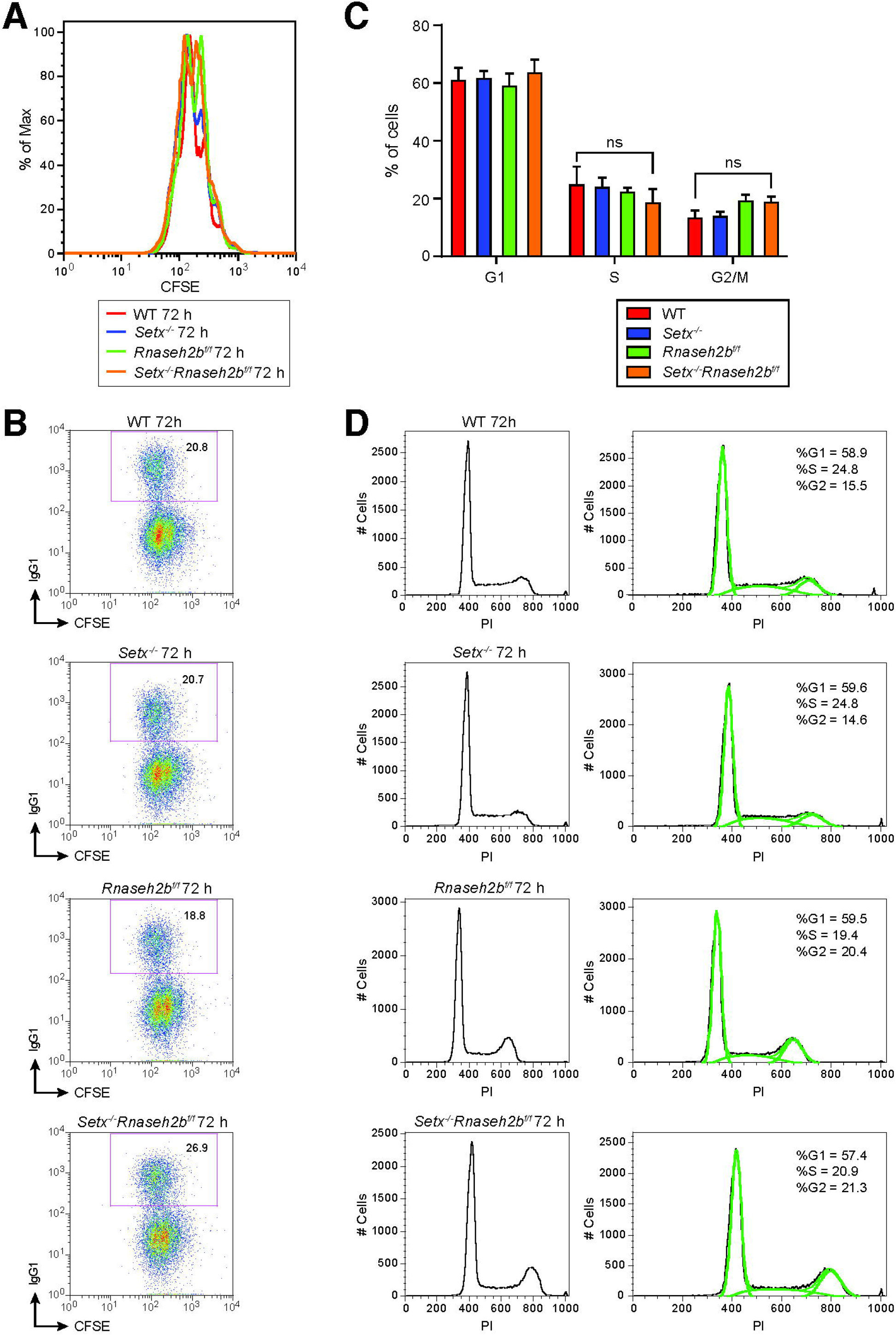
B cell proliferation and cell cycle in response to stimulation. **A**. Flow cytometry analysis of cell proliferation with CFSE staining. CFSE-labeled primary B cells were stimulated with LPS/IL-4/α-RP105 and cell division was measured by CFSE dye dilution at 72 hours post-stimulation. **B**. Representative flow cytometry analyses of IgG1 staining with CFSE staining in spleen primary B cells in response to LPS/IL-4/α-RP105. **C**. The percentage of cells in each cell cycle. Error bars show the SD, statistical analyses were performed using one-way ANOVA (n = 3 mice per genotype). **D**. Representative cell cycle profiles for WT, *Setx*^*-/-*^, *Rnaseh2b*^*f/f*^ and *Setx*^*-/-*^*Rnaseh2b*^*f/f*^ cells stimulated with LPS/IL-4/α-RP105.

**Figure 5-figure supplement 1.**
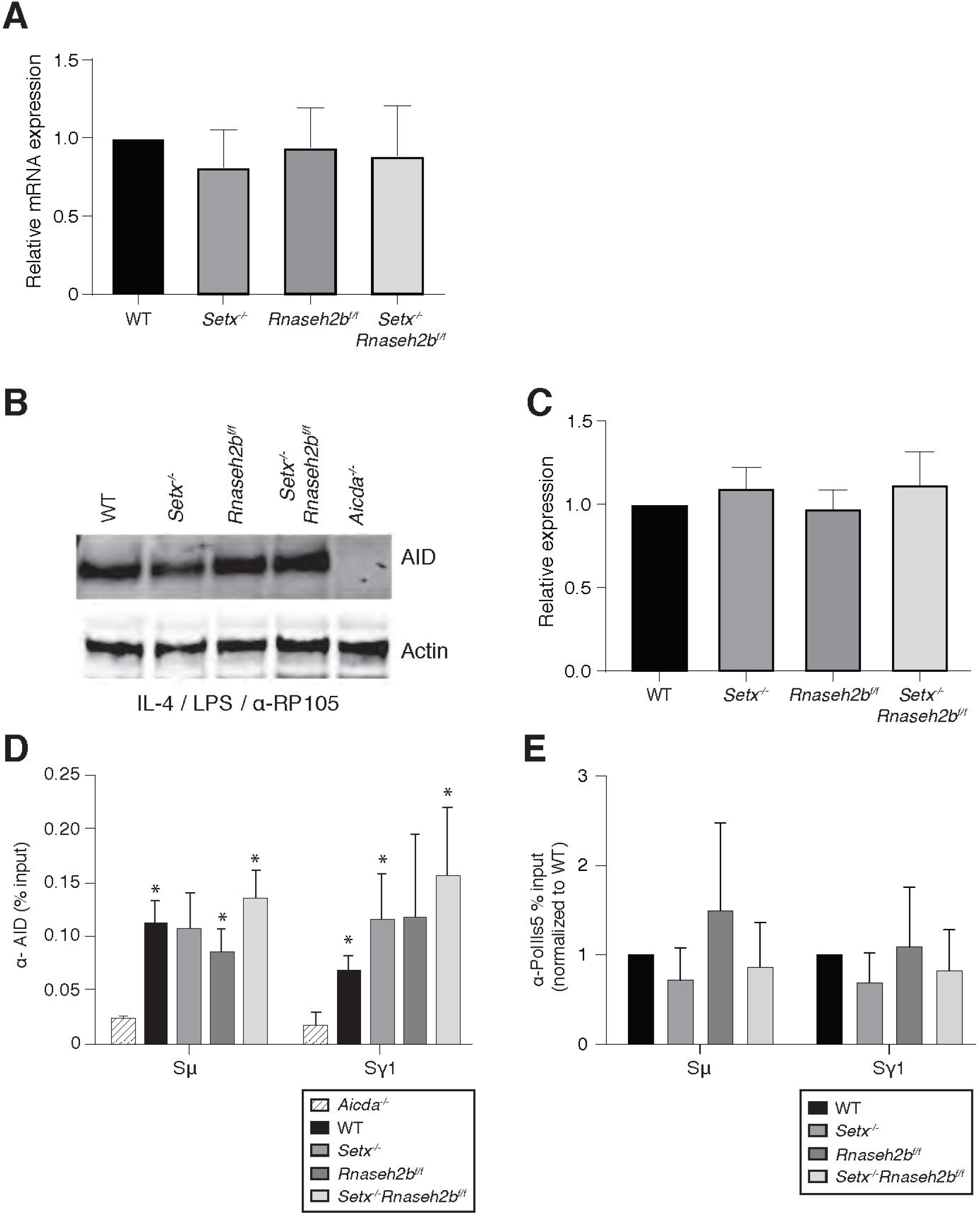
*Aicda* expression, enrichment, and recruitment in cells lacking SETX and RNase H2. **A**. AID mRNA quantitation by RT-qPCR in resting cells, LPS/IL-4/α-RP105, LPS/α-RP105, and α-RP105-only stimulated cells. Error bars show the standard deviation from three independent experiments. **B**. AID protein expression in WT, *Setx*^*-/-*^, *Rnaseh2b*^*f/f*^, *Setx*^*-/-*^*Rnaseh2b*^*f/f*^ and *Aicda*^*-/-*^ cells 72 hours post-stimulation to IgG1. Western blots were probed with an anti-AID antibody and anti-actin as loading control. *Aicda*^*-/-*^ cells were used as a negative control. **C**. Quantification of AID protein expression versus WT related to Figure 5-figure supplement 1B; Error bars show standard deviation; statistical analysis was performed using multiple t-test (n = 3 mice/genotype). **D**. ChIP analysis for AID occupancy in Sμ and Sγ regions of primary B cells in response to LPS/IL-4/α-RP105 stimulation. Relative enrichment was calculated as ChIP/input. Error bars show standard deviation (n = 3 mice/genotype); statistical analysis versus *Aicda*^*-/-*^ control was performed using the students t-test, * denotes p-value < 0.05 compared to *Aicda*^*-/*-^. **E**. ChIP analysis for Pol ll (Ser5) occupancy in Sμ and Sγ regions of primary B cells in response to LPS/IL-4/α-RP105 stimulation. Relative enrichment was calculated as fold change relative to WT set to 1; error bars show standard deviation (n = 5 mice/genotype).

**Figure 6-figure supplement 1.**
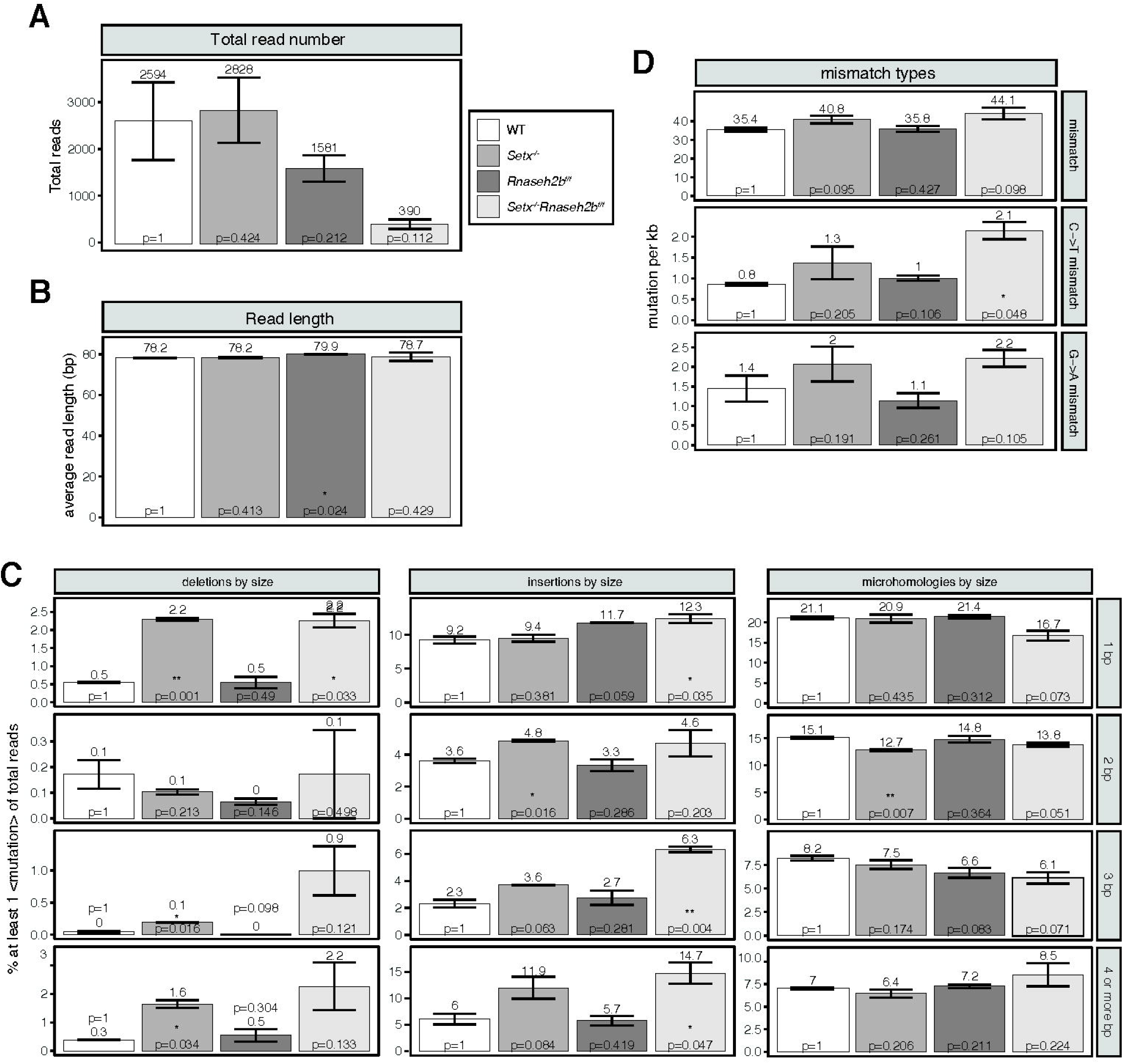
Mutation features of Sμ-Sγ1 junction sites. **A**. Average number of reads from two runs of LAM-HTGTS spanning IgM-IgG1 junctions from WT, *Setx*^*-/-*^, *Rnaseh2b*^*f/f*^, and *Setx*^*-/-*^*Rnaseh2b*^*f/f*^ cells 72 hours post-stimulation with LPs/IL4/α-RP105. **B**. Average read lengths for analyzed junctions spanning IgM-IgG1. **C**. Percentage of reads with different sizes of deletions, insertions or microhomology events at junction sites from WT, *Setx*^*-/-*^, *Rnaseh2b*^*f/f*^, and *Setx*^*-/-*^*Rnaseh2b*^*f/f*^ cells. with 1 bp, 2 bp, 3 bp, and 4 or more bp of each mutation type are shown. **D**. Frequency of mismatches for each genotype within IgM-IgG1 junction reads. Graphs from top to bottom show the total mismatches per kilobase (kb), the number of C to T mismatches per kb, and the number of G to A mismatches per kb. P values are calculated using student’s t-test.

## References

Aguilera, A., & Garcia-Muse, T. (2012). R loops: from transcription byproducts to threats to genome stability. Mol Cell, 46(2), 115–124. doi:10.1016/j.molcel.2012.04.009

Alt, F. W., Zhang, Y., Meng, F. L., Guo, C., & Schwer, B. (2013). Mechanisms of programmed DNA lesions and genomic instability in the immune system. Cell, 152(3), 417–429. doi:10.1016/j.cell.2013.01.007

Bardwell, P. D., Woo, C. J., Wei, K., Li, Z., Martin, A., Sack, S. Z., … Scharff, M. D. (2004). Altered somatic hypermutation and reduced class-switch recombination in exonuclease 1-mutant mice. Nat Immunol, 5(2), 224–229. doi:10.1038/ni1031

Barlow, J. H., Faryabi, R. B., Callen, E., Wong, N., Malhowski, A., Chen, H. T., … Nussenzweig, A. (2013). Identification of early replicating fragile sites that contribute to genome instability. Cell, 152(3), 620–632. doi:10.1016/j.cell.2013.01.006

Basu, U., Meng, F. L., Keim, C., Grinstein, V., Pefanis, E., Eccleston, J., … Alt, F. W. (2011). The RNA exosome targets the AID cytidine deaminase to both strands of transcribed duplex DNA substrates. Cell, 144(3), 353–363. doi:10.1016/j.cell.2011.01.001

Becherel, O. J., Yeo, A. J., Stellati, A., Heng, E. Y., Luff, J., Suraweera, A. M., … Lavin, M. F. (2013). Senataxin plays an essential role with DNA damage response proteins in meiotic recombination and gene silencing. PLoS Genet, 9(4), e1003435. doi:10.1371/journal.pgen.1003435

Bennardo, N., Cheng, A., Huang, N., & Stark, J. M. (2008). Alternative-NHEJ Is a Mechanistically Distinct Pathway of Mammalian Chromosome Break Repair. PLoS Genetics, 4(6). Retrieved from <Go to ISI>://WOS:000260410300010

Boboila, C., Yan, C., Wesemann, D. R., Jankovic, M., Wang, J. H., Manis, J., … Alt, F. W. (2010). Alternative end-joining catalyzes class switch recombination in the absence of both Ku70 and DNA ligase 4. J Exp Med, 207(2), 417–427. doi:10.1084/jem.20092449

Callen, E., Jankovic, M., Difilippantonio, S., Daniel, J. A., Chen, H. T., Celeste, A., … Nussenzweig, A. (2007). ATM prevents the persistence and propagation of chromosome breaks in lymphocytes. Cell, 130(1), 63–75. doi:10.1016/j.cell.2007.06.016

Casellas, R., Nussenzweig, A., Wuerffel, R., Pelanda, R., Reichlin, A., Suh, H., … Nussenzweig, M. C. (1998). Ku80 is required for immunoglobulin isotype switching. EMBO J, 17(8), 2404–2411. doi:10.1093/emboj/17.8.2404

Chang, H. H., Watanabe, G., Gerodimos, C. A., Ochi, T., Blundell, T. L., Jackson, S. P., & Lieber, M. R. (2016). Different DNA End Configurations Dictate Which NHEJ Components Are Most Important for Joining Efficiency. J Biol Chem, 291(47), 24377–24389. doi:10.1074/jbc.M116.752329

Chang, H. H. Y., Pannunzio, N. R., Adachi, N., & Lieber, M. R. (2017). Non-homologous DNA end joining and alternative pathways to double-strand break repair. Nat Rev Mol Cell Biol, 18(8), 495–506. doi:10.1038/nrm.2017.48

Chaudhuri, J., Khuong, C., & Alt, F. W. (2004). Replication protein A interacts with AID to promote deamination of somatic hypermutation targets. Nature, 430(7003), 992–998. doi:10.1038/nature02821

Chaudhuri, J., Tian, M., Khuong, C., Chua, K., Pinaud, E., & Alt, F. W. (2003). Transcription-targeted DNA deamination by the AID antibody diversification enzyme. Nature, 422(6933), 726–730. doi:10.1038/nature01574

Chon, H., Sparks, J. L., Rychlik, M., Nowotny, M., Burgers, P. M., Crouch, R. J., & Cerritelli, S. M. (2013). RNase H2 roles in genome integrity revealed by unlinking its activities. Nucleic Acids Res, 41(5), 3130–3143. doi:10.1093/nar/gkt027

Cohen, S., Puget, N., Lin, Y. L., Clouaire, T., Aguirrebengoa, M., Rocher, V., … Legube, G. (2018). Senataxin resolves RNA:DNA hybrids forming at DNA double-strand breaks to prevent translocations. Nat Commun, 9(1), 533. doi:10.1038/s41467-018-02894-w

Costantino, L., & Koshland, D. (2018). Genome-wide Map of R-Loop-Induced Damage Reveals How a Subset of R-Loops Contributes to Genomic Instability. Mol Cell, 71(4), 487–497 e483. doi:10.1016/j.molcel.2018.06.037

Crossley, M. P., Bocek, M., & Cimprich, K. A. (2019). R-Loops as Cellular Regulators and Genomic Threats. Mol Cell, 73(3), 398–411. doi:10.1016/j.molcel.2019.01.024

Crowe, J. L., Shao, Z., Wang, X. S., Wei, P. C., Jiang, W., Lee, B. J., … Zha, S. (2018). Kinase-dependent structural role of DNA-PKcs during immunoglobulin class switch recombination. Proc Natl Acad Sci U S A. doi:10.1073/pnas.1808490115

Daniels, G. A., & Lieber, M. R. (1995). RNA:DNA complex formation upon transcription of immunoglobulin switch regions: implications for the mechanism and regulation of class switch recombination. Nucleic Acids Res, 23(24), 5006–5011. Retrieved from https://www.ncbi.nlm.nih.gov/pubmed/8559658

Du, L., van der Burg, M., Popov, S. W., Kotnis, A., van Dongen, J. J., Gennery, A. R., & Pan-Hammarstrom, Q. (2008). Involvement of Artemis in nonhomologous end-joining during immunoglobulin class switch recombination. J Exp Med, 205(13), 3031–3040. doi:10.1084/jem.20081915

Franco, S., Murphy, M. M., Li, G., Borjeson, T., Boboila, C., & Alt, F. W. (2008). DNA-PKcs and Artemis function in the end-joining phase of immunoglobulin heavy chain class switch recombination. Journal of Experimental Medicine, 205(3), 557–564. doi:10.1084/jem.20080044

Ginno, P. A., Lott, P. L., Christensen, H. C., Korf, I., & Chedin, F. (2012). R-loop formation is a distinctive characteristic of unmethylated human CpG island promoters. Mol Cell, 45(6), 814–825. doi:10.1016/j.molcel.2012.01.017

Grigera, F., Wuerffel, R., & Kenter, A. L. (2017). MBD4 Facilitates Immunoglobulin Class Switch Recombination. Mol Cell Biol, 37(2). doi:10.1128/MCB.00316-16

Gu, Y., Seidl, K. J., Rathbun, G. A., Zhu, C., Manis, J. P., van der Stoep, N., … Alt, F. W. (1997). Growth retardation and leaky SCID phenotype of Ku70-deficient mice. Immunity, 7(5), 653–665. Retrieved from https://www.ncbi.nlm.nih.gov/pubmed/9390689

Hein, K., Lorenz, M. G., Siebenkotten, G., Petry, K., Christine, R., & Radbruch, A. (1998). Processing of switch transcripts is required for targeting of antibody class switch recombination. J Exp Med, 188(12), 2369–2374. doi:10.1084/jem.188.12.2369

Hiller, B., Achleitner, M., Glage, S., Naumann, R., Behrendt, R., & Roers, A. (2012). Mammalian RNase H2 removes ribonucleotides from DNA to maintain genome integrity. J Exp Med, 209(8), 1419–1426. doi:10.1084/jem.20120876

Hodgkin, P. D., Lee, J. H., & Lyons, A. B. (1996). B cell differentiation and isotype switching is related to division cycle number. J Exp Med, 184(1), 277–281. doi:10.1084/jem.184.1.277

Hodroj, D., Recolin, B., Serhal, K., Martinez, S., Tsanov, N., Abou Merhi, R., & Maiorano, D. (2017). An ATR-dependent function for the Ddx19 RNA helicase in nuclear R-loop metabolism. EMBO J, 36(9), 1182–1198. doi:10.15252/embj.201695131

Hu, J., Meyers, R. M., Dong, J., Panchakshari, R. A., Alt, F. W., & Frock, R. L. (2016). Detecting DNA double-stranded breaks in mammalian genomes by linear amplification-mediated high-throughput genome-wide translocation sequencing. Nat Protoc, 11(5), 853–871. doi:10.1038/nprot.2016.043

Huang, F. T., Yu, K., Balter, B. B., Selsing, E., Oruc, Z., Khamlichi, A. A., … Lieber, M. R. (2007). Sequence dependence of chromosomal R-loops at the immunoglobulin heavy-chain Smu class switch region. Mol Cell Biol, 27(16), 5921–5932. doi:10.1128/MCB.00702-07

Huang, S. N., Williams, J. S., Arana, M. E., Kunkel, T. A., & Pommier, Y. (2017). Topoisomerase I-mediated cleavage at unrepaired ribonucleotides generates DNA double-strand breaks. EMBO J, 36(3), 361–373. doi:10.15252/embj.201592426

Kazadi, D., Lim, J., Rothschild, G., Grinstein, V., Laffleur, B., Becherel, O., … Basu, U. (2020). Effects of senataxin and RNA exosome on B-cell chromosomal integrity. Heliyon, 6(3), e03442. doi:10.1016/j.heliyon.2020.e03442

Keskin, H., Shen, Y., Huang, F., Patel, M., Yang, T., Ashley, K., … Storici, F. (2014). Transcript-RNA-templated DNA recombination and repair. Nature, 515(7527), 436–439. doi:10.1038/nature13682

Kuznetsov, V. A., Bondarenko, V., Wongsurawat, T., Yenamandra, S. P., & Jenjaroenpun, P. (2018). Toward predictive R-loop computational biology: genome-scale prediction of R-loops reveals their association with complex promoter structures, G-quadruplexes and transcriptionally active enhancers (vol 46, pg 7566, 2018). Nucleic Acids Research, 46(15), 8023–8023. Retrieved from <Go to ISI>://WOS:000444148100047 https://www.ncbi.nlm.nih.gov/pmc/articles/PMC6125685/pdf/gky690.pdf

Lee-Theilen, M., Matthews, A. J., Kelly, D., Zheng, S., & Chaudhuri, J. (2011). CtIP promotes microhomology-mediated alternative end joining during class-switch recombination. Nat Struct Mol Biol, 18(1), 75–79. doi:10.1038/nsmb.1942

Li, L., Halaby, M. J., Hakem, A., Cardoso, R., El Ghamrasni, S., Harding, S., … Hakem, R. (2010). Rnf8 deficiency impairs class switch recombination, spermatogenesis, and genomic integrity and predisposes for cancer. J Exp Med, 207(5), 983–997. doi:10.1084/jem.20092437

Lim, G., & Hohng, S. (2020). Single-molecule fluorescence studies on cotranscriptional G-quadruplex formation coupled with R-loop formation. Nucleic Acids Res, 48(16), 9195–9203. doi:10.1093/nar/gkaa695

Liu, D., Goodman, M. F., Pham, P., Yu, K., Hsieh, C. L., & Lieber, M. R. (2021). The mRNA tether model for activation-induced deaminase and its relevance for Ig somatic hypermutation and class switch recombination. DNA Repair (Amst), 110, 103271. doi:10.1016/j.dnarep.2021.103271

Lumsden, J. M., McCarty, T., Petiniot, L. K., Shen, R., Barlow, C., Wynn, T. A., … Hodes, R. J. (2004). Immunoglobulin class switch recombination is impaired in Atm-deficient mice. Journal of Experimental Medicine, 200(9), 1111–1121. Retrieved from <Go to ISI>://WOS:000224961600004

Lyons, A. B., & Parish, C. R. (1994). Determination of lymphocyte division by flow cytometry. J Immunol Methods, 171(1), 131–137. Retrieved from https://www.ncbi.nlm.nih.gov/pubmed/8176234

Manis, J. P., Dudley, D., Kaylor, L., & Alt, F. W. (2002). IgH class switch recombination to IgG1 in DNA-PKcs-deficient B cells. Immunity, 16(4), 607–617. doi:10.1016/s1074-7613(02)00306-0

Marchalot, A., Ashi, M. O., Lambert, J. M., Carrion, C., Lecardeur, S., Srour, N., … Le Pennec, S. (2020). Uncoupling Splicing From Transcription Using Antisense Oligonucleotides Reveals a Dual Role for I Exon Donor Splice Sites in Antibody Class Switching. Front Immunol, 11, 780. doi:10.3389/fimmu.2020.00780

Marnef, A., & Legube, G. (2021). R-loops as Janus-faced modulators of DNA repair. Nat Cell Biol, 23(4), 305–313. doi:10.1038/s41556-021-00663-4

Mateos-Gomez, P. A., Gong, F., Nair, N., Miller, K. M., Lazzerini-Denchi, E., & Sfeir, A. (2015). Mammalian polymerase theta promotes alternative NHEJ and suppresses recombination. Nature, 518(7538), 254–257. doi:10.1038/nature14157

Maul, R. W., Chon, H., Sakhuja, K., Cerritelli, S. M., Gugliotti, L. A., Gearhart, P. J., & Crouch, R. J. (2017). R-Loop Depletion by Over-expressed RNase H1 in Mouse B Cells Increases Activation-Induced Deaminase Access to the Transcribed Strand without Altering Frequency of Isotype Switching. J Mol Biol, 429(21), 3255–3263. doi:10.1016/j.jmb.2016.12.020

Mazina, O. M., Keskin, H., Hanamshet, K., Storici, F., & Mazin, A. V. (2017). Rad52 Inverse Strand Exchange Drives RNA-Templated DNA Double-Strand Break Repair. Mol Cell, 67(1), 19–29 e13. doi:10.1016/j.molcel.2017.05.019

McDevitt, S., Rusanov, T., Kent, T., Chandramouly, G., & Pomerantz, R. T. (2018). How RNA transcripts coordinate DNA recombination and repair. Nat Commun, 9(1), 1091. doi:10.1038/s41467-018-03483-7

Mcdonell, M. W., Simon, M. N., & Studier, F. W. (1977). Analysis of Restriction Fragments of T7 DNA and Determination of Molecular-Weights by Electrophoresis in Neutral and Alkaline Gels. Journal of Molecular Biology, 110(1), 119–146. Retrieved from <Go to ISI>://WOS:A1977DC70700008

Muramatsu, M., Kinoshita, K., Fagarasan, S., Yamada, S., Shinkai, Y., & Honjo, T. (2000). Class switch recombination and hypermutation require activation-induced cytidine deaminase (AID), a potential RNA editing enzyme. Cell, 102(5), 553–563. Retrieved from https://www.ncbi.nlm.nih.gov/pubmed/11007474

Nick McElhinny, S. A., Kumar, D., Clark, A. B., Watt, D. L., Watts, B. E., Lundstrom, E. B., … Kunkel, T. A. (2010). Genome instability due to ribonucleotide incorporation into DNA. Nat Chem Biol, 6(10), 774–781. doi:10.1038/nchembio.424

Nowak, U., Matthews, A. J., Zheng, S., & Chaudhuri, J. (2011). The splicing regulator PTBP2 interacts with the cytidine deaminase AID and promotes binding of AID to switch-region DNA. Nat Immunol, 12(2), 160–166. doi:10.1038/ni.1977

Nussenzweig, A., & Nussenzweig, M. C. (2007). A backup DNA repair pathway moves to the forefront. Cell, 131(2), 223–225. doi:10.1016/j.cell.2007.10.005

Parsa, J. Y., Ramachandran, S., Zaheen, A., Nepal, R. M., Kapelnikov, A., Belcheva, A., … Martin, A. (2012). Negative supercoiling creates single-stranded patches of DNA that are substrates for AID-mediated mutagenesis. PLoS Genet, 8(2), e1002518. doi:10.1371/journal.pgen.1002518

Pavri, R., Gazumyan, A., Jankovic, M., Di Virgilio, M., Klein, I., Ansarah-Sobrinho, C., … Nussenzweig, M. C. (2010). Activation-induced cytidine deaminase targets DNA at sites of RNA polymerase II stalling by interaction with Spt5. Cell, 143(1), 122–133. doi:10.1016/j.cell.2010.09.017

Pham, P., Bransteitter, R., Petruska, J., & Goodman, M. F. (2003). Processive AID-catalysed cytosine deamination on single-stranded DNA simulates somatic hypermutation. Nature, 424(6944), 103–107. doi:10.1038/nature01760

Prado, F., & Aguilera, A. (2005). Impairment of replication fork progression mediates RNA polII transcription-associated recombination. EMBO J, 24(6), 1267–1276. doi:10.1038/sj.emboj.7600602

Qiao, Q., Wang, L., Meng, F. L., Hwang, J. K., Alt, F. W., & Wu, H. (2017). AID Recognizes Structured DNA for Class Switch Recombination. Mol Cell, 67(3), 361–373 e364. doi:10.1016/j.molcel.2017.06.034

Ramiro, A. R., Jankovic, M., Eisenreich, T., Difilippantonio, S., Chen-Kiang, S., Muramatsu, M., … Nussenzweig, M. C. (2004). AID is required for c-myc/IgH chromosome translocations in vivo. Cell, 118(4), 431–438. doi:10.1016/j.cell.2004.08.006

Ramiro, A. R., Stavropoulos, P., Jankovic, M., & Nussenzweig, M. C. (2003a). Transcription enhances AID-mediated cytidine deamination by exposing single-stranded DNA on the nontemplate strand. Nat Immunol, 4(5), 452–456. doi:10.1038/ni920

Ramiro, A. R., Stavropoulos, P., Jankovic, M., & Nussenzweig, M. C. (2003b). Transcription enhances AID-mediated cytidine deamination by exposing single-stranded DNA on the nontemplate strand. Nature Immunology, 4(5), 452–456. Retrieved from <Go to ISI>://WOS:000182665400012 https://www.nature.com/articles/ni920.pdf

Reaban, M. E., & Griffin, J. A. (1990). Induction of RNA-stabilized DNA conformers by transcription of an immunoglobulin switch region. Nature, 348(6299), 342–344. doi:10.1038/348342a0

Reijns, M. A., Rabe, B., Rigby, R. E., Mill, P., Astell, K. R., Lettice, L. A., … Jackson, A. P. (2012). Enzymatic removal of ribonucleotides from DNA is essential for mammalian genome integrity and development. Cell, 149(5), 1008–1022. doi:10.1016/j.cell.2012.04.011

Reina-San-Martin, B., Chen, H. T., Nussenzweig, A., & Nussenzweig, M. C. (2004). ATM is required for efficient recombination between immunoglobulin switch regions. Journal of Experimental Medicine, 200(9), 1103–1110. Retrieved from <Go to ISI>://WOS:000224961600003

Revy, P., Muto, T., Levy, Y., Geissmann, F., Plebani, A., Sanal, O., … Durandy, A. (2000). Activation-induced cytidine deaminase (AID) deficiency causes the autosomal recessive form of the Hyper-IgM syndrome (HIGM2). Cell, 102(5), 565–575. doi:10.1016/s0092-8674(00)00079-9

Ribeiro de Almeida, C., Dhir, S., Dhir, A., Moghaddam, A. E., Sattentau, Q., Meinhart, A., & Proudfoot, N. J. (2018). RNA Helicase DDX1 Converts RNA G-Quadruplex Structures into R-Loops to Promote IgH Class Switch Recombination. Mol Cell, 70(4), 650–662 e658. doi:10.1016/j.molcel.2018.04.001

Richard, P., Feng, S., Tsai, Y. L., Li, W., Rinchetti, P., Muhith, U., … Manley, J. L. (2020). SETX (senataxin), the helicase mutated in AOA2 and ALS4, functions in autophagy regulation. Autophagy, 1–18. doi:10.1080/15548627.2020.1796292

Rickert, R. C., Roes, J., & Rajewsky, K. (1997). B lymphocyte-specific, Cre-mediated mutagenesis in mice. Nucleic Acids Res, 25(6), 1317–1318. Retrieved from https://www.ncbi.nlm.nih.gov/pubmed/9092650

Rivera-Munoz, P., Soulas-Sprauel, P., Le Guyader, G., Abramowski, V., Bruneau, S., Fischer, A., … de Villartay, J. P. (2009). Reduced immunoglobulin class switch recombination in the absence of Artemis. Blood, 114(17), 3601–3609. doi:10.1182/blood-2008-11-188383

Robbiani, D. F., Bothmer, A., Callen, E., Reina-San-Martin, B., Dorsett, Y., Difilippantonio, S., … Nussenzweig, M. C. (2008). AID is required for the chromosomal breaks in c-myc that lead to c-myc/IgH translocations. Cell, 135(6), 1028–1038. doi:10.1016/j.cell.2008.09.062

Robbiani, D. F., Bunting, S., Feldhahn, N., Bothmer, A., Camps, J., Deroubaix, S., … Nussenzweig, M. C. (2009). AID produces DNA double-strand breaks in non-Ig genes and mature B cell lymphomas with reciprocal chromosome translocations. Mol Cell, 36(4), 631–641. doi:10.1016/j.molcel.2009.11.007

Roberts, S. A., Strande, N., Burkhalter, M. D., Strom, C., Havener, J. M., Hasty, P., & Ramsden, D. A. (2010). Ku is a 5’-dRP/AP lyase that excises nucleotide damage near broken ends. Nature, 464(7292), 1214–1217. doi:10.1038/nature08926

Roy, D., Yu, K., & Lieber, M. R. (2008). Mechanism of R-loop formation at immunoglobulin class switch sequences. Mol Cell Biol, 28(1), 50–60. doi:10.1128/MCB.01251-07

Santos, M. A., Huen, M. S., Jankovic, M., Chen, H. T., Lopez-Contreras, A. J., Klein, I. A., … Nussenzweig, A. (2010). Class switching and meiotic defects in mice lacking the E3 ubiquitin ligase RNF8. J Exp Med, 207(5), 973–981. doi:10.1084/jem.20092308

Sanz, L. A., Hartono, S. R., Lim, Y. W., Steyaert, S., Rajpurkar, A., Ginno, P. A., … Chedin, F. (2016). Prevalent, Dynamic, and Conserved R-Loop Structures Associate with Specific Epigenomic Signatures in Mammals. Mol Cell, 63(1), 167–178. doi:10.1016/j.molcel.2016.05.032

Schenten, D., Kracker, S., Esposito, G., Franco, S., Klein, U., Murphy, M., … Rajewsky, K. (2009). Pol zeta ablation in B cells impairs the germinal center reaction, class switch recombination, DNA break repair, and genome stability. J Exp Med, 206(2), 477–490. doi:10.1084/jem.20080669

Schrader, C. E., Vardo, J., & Stavnezer, J. (2002). Role for mismatch repair proteins Msh2, Mlh1, and Pms2 in immunoglobulin class switching shown by sequence analysis of recombination junctions. J Exp Med, 195(3), 367–373. Retrieved from https://www.ncbi.nlm.nih.gov/pubmed/11828012

Serrano-Benitez, A., Cortes-Ledesma, F., & Ruiz, J. F. (2019). “An End to a Means”: How DNA-End Structure Shapes the Double-Strand Break Repair Process. Front Mol Biosci, 6, 153. doi:10.3389/fmolb.2019.00153

So, C. C., & Martin, A. (2019). DSB structure impacts DNA recombination leading to class switching and chromosomal translocations in human B cells. PLoS Genet, 15(4), e1008101. doi:10.1371/journal.pgen.1008101

Sohail, A., Klapacz, J., Samaranayake, M., Ullah, A., & Bhagwat, A. S. (2003). Human activation-induced cytidine deaminase causes transcription-dependent, strand-biased C to U deaminations. Nucleic Acids Research, 31(12), 2990–2994. Retrieved from <Go to ISI>://WOS:000183832000002 https://www.ncbi.nlm.nih.gov/pmc/articles/PMC162340/pdf/gkg464.pdf

Sollier, J., Stork, C. T., Garcia-Rubio, M. L., Paulsen, R. D., Aguilera, A., & Cimprich, K. A. (2014). Transcription-coupled nucleotide excision repair factors promote R-loop-induced genome instability. Mol Cell, 56(6), 777–785. doi:10.1016/j.molcel.2014.10.020

Song, C. L., Hotz-Wagenblatt, A., Voit, R., & Grummt, I. (2017). SIRT7 and the DEAD-box helicase DDX21 cooperate to resolve genomic R loops and safeguard genome stability. Genes & Development, 31(13), 1370–1381. Retrieved from <Go to ISI>://WOS:000407611300007

Soulas-Sprauel, P., Le Guyader, G., Rivera-Munoz, P., Abramowski, V., Olivier-Martin, C., Goujet-Zalc, C., … de Villartay, J. P. (2007). Role for DNA repair factor XRCC4 in immunoglobulin class switch recombination. Journal of Experimental Medicine, 204(7), 1717–1727. Retrieved from <Go to ISI>://WOS:000247998000022 https://www.ncbi.nlm.nih.gov/pmc/articles/PMC2118634/pdf/jem2041717.pdf

Stanlie, A., Begum, N. A., Akiyama, H., & Honjo, T. (2012). The DSIF subunits Spt4 and Spt5 have distinct roles at various phases of immunoglobulin class switch recombination. PLoS Genet, 8(4), e1002675. doi:10.1371/journal.pgen.1002675

Stavnezer, J., & Schrader, C. E. (2014). IgH chain class switch recombination: mechanism and regulation. J Immunol, 193(11), 5370–5378. doi:10.4049/jimmunol.1401849

Stolz, R., Sulthana, S., Hartono, S. R., Malig, M., Benham, C. J., & Chedin, F. (2019). Interplay between DNA sequence and negative superhelicity drives R-loop structures. Proceedings of the National Academy of Sciences of the United States of America, 116(13), 6260–6269. Retrieved from <Go to ISI>://WOS:000462382800068 https://escholarship.org/content/qt0vz4t8b4/qt0vz4t8b4.pdf?t=pzewzl

Storici, F., Bebenek, K., Kunkel, T. A., Gordenin, D. A., & Resnick, M. A. (2007). RNA-templated DNA repair. Nature, 447(7142), 338–341. doi:10.1038/nature05720

Stork, C. T., Bocek, M., Crossley, M. P., Sollier, J., Sanz, L. A., Chedin, F., … Cimprich, K. A. (2016). Co-transcriptional R-loops are the main cause of estrogen-induced DNA damage. Elife, 5. doi:10.7554/eLife.17548

Strande, N., Roberts, S. A., Oh, S., Hendrickson, E. A., & Ramsden, D. A. (2012). Specificity of the dRP/AP lyase of Ku promotes nonhomologous end joining (NHEJ) fidelity at damaged ends. J Biol Chem, 287(17), 13686–13693. doi:10.1074/jbc.M111.329730

Symington, L. S. (2016). Mechanism and regulation of DNA end resection in eukaryotes. Crit Rev Biochem Mol Biol, 51(3), 195–212. doi:10.3109/10409238.2016.1172552

Uchimura, Y., Barton, L. F., Rada, C., & Neuberger, M. S. (2011). REG-gamma associates with and modulates the abundance of nuclear activation-induced deaminase. J Exp Med, 208(12), 2385–2391. doi:10.1084/jem.20110856

Waisertreiger, I., Popovich, K., Block, M., Anderson, K. R., & Barlow, J. H. (2020). Visualizing locus-specific sister chromatid exchange reveals differential patterns of replication stress-induced fragile site breakage. Oncogene, 39(6), 1260–1272. doi:10.1038/s41388-019-1054-5

Ward, I. M., Reina-San-Martin, B., Olaru, A., Minn, K., Tamada, K., Lau, J. S., … Chen, J. (2004). 53BP1 is required for class switch recombination. J Cell Biol, 165(4), 459–464. doi:10.1083/jcb.200403021

Wiedemann, E. M., Peycheva, M., & Pavri, R. (2016). DNA Replication Origins in Immunoglobulin Switch Regions Regulate Class Switch Recombination in an R-Loop-Dependent Manner. Cell Rep, 17(11), 2927–2942. doi:10.1016/j.celrep.2016.11.041

Williams, J. S., & Kunkel, T. A. (2014). Ribonucleotides in DNA: origins, repair and consequences. DNA Repair (Amst), 19, 27–37. doi:10.1016/j.dnarep.2014.03.029

Williams, J. S., Smith, D. J., Marjavaara, L., Lujan, S. A., Chabes, A., & Kunkel, T. A. (2013). Topoisomerase 1-mediated removal of ribonucleotides from nascent leading-strand DNA. Mol Cell, 49(5), 1010–1015. doi:10.1016/j.molcel.2012.12.021

Willmann, K. L., Milosevic, S., Pauklin, S., Schmitz, K. M., Rangam, G., Simon, M. T., … Petersen-Mahrt, S. K. (2012). A role for the RNA pol II-associated PAF complex in AID-induced immune diversification. J Exp Med, 209(11), 2099–2111. doi:10.1084/jem.20112145

Yamane, A., Resch, W., Kuo, N., Kuchen, S., Li, Z., Sun, H. W., … Casellas, R. (2011). Deep-sequencing identification of the genomic targets of the cytidine deaminase AID and its cofactor RPA in B lymphocytes. Nat Immunol, 12(1), 62–69. doi:10.1038/ni.1964

Yan, C. T., Boboila, C., Souza, E. K., Franco, S., Hickernell, T. R., Murphy, M., … Alt, F. W. (2007). IgH class switching and translocations use a robust non-classical end-joining pathway. Nature, 449(7161), 478–482. doi:10.1038/nature06020

Yin, J., Liu, M., Liu, Y., & Hu, J. (2019). Improved HTGTS for CRISPR/Cas9 off-target detection. Bio Protoc, 9(9), e3229. doi:10.21769/BioProtoc.3229

Yin, J., Liu, M., Liu, Y., Wu, J., Gan, T., Zhang, W., … Hu, J. (2019). Optimizing genome editing strategy by primer-extension-mediated sequencing. Cell Discov, 5, 18. doi:10.1038/s41421-019-0088-8

Yousefzadeh, M. J., Wyatt, D. W., Takata, K., Mu, Y., Hensley, S. C., Tomida, J., … Wood, R. D. (2014). Mechanism of suppression of chromosomal instability by DNA polymerase POLQ. PLoS Genet, 10(10), e1004654. doi:10.1371/journal.pgen.1004654

Yu, A. M., & McVey, M. (2010). Synthesis-dependent microhomology-mediated end joining accounts for multiple types of repair junctions. Nucleic Acids Res, 38(17), 5706–5717. doi:10.1093/nar/gkq379

Yu, K., Chedin, F., Hsieh, C. L., Wilson, T. E., & Lieber, M. R. (2003). R-loops at immunoglobulin class switch regions in the chromosomes of stimulated B cells. Nat Immunol, 4(5), 442–451. doi:10.1038/ni919

Zan, H., Tat, C., Qiu, Z., Taylor, J. R., Guerrero, J. A., Shen, T., & Casali, P. (2017). Rad52 competes with Ku70/Ku86 for binding to S-region DSB ends to modulate antibody class-switch DNA recombination. Nat Commun, 8, 14244. doi:10.1038/ncomms14244

Zhang, Z. Z., Pannunzio, N. R., Hsieh, C. L., Yu, K., & Lieber, M. R. (2014). The role of G-density in switch region repeats for immunoglobulin class switch recombination. Nucleic Acids Res, 42(21), 13186–13193. doi:10.1093/nar/gku1100

Zheng, S. M., Vuong, B. Q., Vaidyanathan, B., Lin, J. Y., Huang, F. T., & Chaudhuri, J. (2015). Non-coding RNA Generated following Lariat Debranching Mediates Targeting of AID to DNA. Cell, 161(4), 762–773. doi:10.1016/j.cell.2015.03.020

